# Strain-specific higher-order interactions break phylogenetic conservatism in microbial communities

**DOI:** 10.64898/2026.10.10.758215

**Authors:** Cordero-Martínez Emmanuel, Navarro-Miranda Marisol, Ramírez-Sánchez Obed, Edder D. Bustos-Díaz, Higareda-Alvear Víctor Manuel, Gabriel Moreno-Hagelsieb, Rocha-Jorge, Hernández-Rosales Maribel, Olmedo-Álvarez Gabriela

## Abstract

Whether strains of the same species sharing the same ecological role are truly interchangeable has not been directly tested experimentally in communities, where higher-order interactions govern outcomes. Using 79 co-occurring Bacillaceae isolates from the Cuatrociénegas Basin, we combined whole-genome analyses with systematic strain substitutions in BARS, a three-species community in which a resistant strain protects a sensitive strain from an antagonist. Trait conservatism strongly predicted pairwise antagonistic outcomes. Yet across 34 assembled triads, 13 showed rescue beyond pairwise expectations and five showed suppression; single-strain substitutions rarely restored protection, although protection was recovered in one fully reconfigured triad. These outcomes could not be inferred from taxonomic relatedness, pangenomic content or pairwise assays; phylogenetic conservatism thus reaches its limit at the transition from pairwise to higher-order interactions. To explain this limit, we propose that strains behave as non-equivalent biological tiles whose interaction edges must align, a *dynamic biological tessellation*.

## Introduction

The ability to understand how the evolutionary information encoded within a cell dictates its ecological niche, and whether this genomic heritage is sufficient to govern multi-species assembly, remains a fundamental quest in microbial ecology. Microbial communities are structured by complex interaction networks shaped by a diverse chemical arsenal, including antibiotics, bacteriocins, and other biosynthetic gene clusters (BGCs), that determine ecosystem stability and function(Hibbing *et al.*, 2010; Faust & Raes, 2012; Coyte *et al.*, 2015; Ghoul & Mitri, 2016). Comparative genomics has linked BGC repertoires and metabolic capacity to phylogenetic lineage (Russel *et al.*, 2017; Xia *et al.*, 2022), leading to the widespread but rarely tested assumption that closely related strains are functionally redundant and interchangeable within their local habitats (Losos, 2008; Wiens *et al.*, 2010). However, the assumption that ecologically equivalent conspecifics are truly substitutable has not been directly tested in a community context, where emergent higher-order interactions ultimately dictate ecological outcomes.

Phylogenetic trait conservatism provides the conceptual backbone for predicting ecological roles from genomic content. Genetic and physiological constraints limit the rate at which traits diverge, such that closely related organisms tend to retain similar phenotypes and perform comparable ecological functions, a principle with direct consequences for functional redundancy and interchangeability (Louca *et al.*, 2018; Jia & Whalen, 2020). At the community level, this logic extends to community conservatism: the tendency of phylogenetically related microorganisms to co-occur in compositionally similar communities, a pattern confirmed globally across all major microbial phyla (Malfertheiner *et al.*, 2026) and locally in our system, where lineage constrains physiological traits more strongly than environment of origin (Hurtado-Bautista *et al.*, 2021). However, the predictive power of phylogeny declines as trait complexity increases (Martiny *et al.*, 2015). Although strains sharing more than 95% Average Nucleotide Identity (ANI) are commonly classified as conspecifics (Konstantinidis & Tiedje, 2005; Jain *et al.*, 2018), genomic similarity does not guarantee ecological equivalence. Closely related genomes can encode substantial variation in metabolic outputs (Niccum *et al.*, 2020; Patnode *et al.*, 2021; Ashkenazy & Weigel, 2025), and strain-level ecological dynamics can decouple at genetic distances as small as 100 single-nucleotide polymorphisms (Goyal *et al.*, 2022). Consequently, replicated communities assembled under identical conditions diverge at the strain level rather than at higher taxonomic ranks (Leventhal *et al.*, 2018). This microdiversity, defined as functional variation between strains of the same species driven by accessory genome turnover and horizontal gene transfer (Larkin & Martiny, 2017; García-García *et al.*, 2019), creates a barrier to predicting ecological roles from genomic content alone, precisely where such prediction is most needed.

This predictability gap is further compounded by the scale-dependence of microbial interactions. At a pairwise scale, phylogenetic relatedness broadly predicts ecological outcomes. But when interactions extend beyond pairs to include additional species, higher-order interactions (HOIs) emerge, where the presence of a third organism modifies the outcome of pairwise interactions in ways that cannot be inferred from the pair alone (Kleinhesselink *et al.*, 2022). Beyond reversing antagonistic relationships and reshaping competitive hierarchies (Kelsic *et al.*, 2015; Swain *et al.*, 2021; Aguilar-Salinas & Olmedo-Álvarez, 2023), HOIs are increasingly recognized as key determinants of community stability (Bairey *et al.*, 2016; Morin *et al.*, 2022; Chang *et al.*, 2023); theory predicts that they can stabilize otherwise unstable competitive networks (Grilli *et al.*, 2017; Levine *et al.*, 2017). Yet their genomic basis has received comparatively little attention (Sanchez-Gorostiaga *et al.*, 2019; van den Berg *et al.*, 2022; Oña *et al.*, 2025), and it remains unknown whether the capacity to engage in multi-way interactions is a conserved trait governed by ancestral rules or an emergent property unique to specific strain combinations that escape genomic prediction.

Here we directly test whether genomic similarity translates into functional interchangeability from pairwise to community-level interactions. We leverage 79 Bacillaceae isolates from the shallow sediments of El Churince in the Cuatrociénegas Basin, an oligotrophic system harboring microbial communities with deep evolutionary roots and extensive microdiversity (Moreno-Letelier *et al.*, 2012; Souza *et al.*, 2018; González-Sánchez *et al.*, 2026), where phylogenetically related strains consistently co-occur, satisfying the community conservatism condition by design (Malfertheiner *et al.*, 2026).

This collection is also a long-standing model of microbial interaction networks: pairwise antagonism among these isolates structures a hierarchical network (Pérez-Gutiérrez *et al.*, 2013), is sufficient to generate community patches in a computational model (Zapién-Campos *et al.*, 2015), and underlies both emergent protection in a three-species community (Aguilar-Salinas & Olmedo-Álvarez, 2023) and dynamically induced spatial segregation in multispecies suspensions (Gallardo-Navarro *et al.*, 2025). We build on the BARS synthetic community model, a minimal three-species system where a resistant strain protects a sensitive strain from an antagonist (Aguilar-Salinas & Olmedo-Álvarez, 2023). By systematically replacing each member with conspecifics sharing the same ecological role, we linked interaction phenotypes to whole-genome features, including pangenomic architecture and BGC repertoires, to test the phylogenetic reach of conservatism from individual traits to pairwise interactions and, ultimately, to emergent community behavior. We show that trait conservatism is robust at the pairwise scale: competitive capacity and BGC profiles are organized by phylogenetic lineage, even across clades with contrasting levels of accessory genome variation. However, when strains equivalent by genomic, taxonomic, and pairwise-role criteria are assembled into three-species communities, single-strain substitutions fail to restore protection, and triads deviate from null expectations through both emergent rescue and suppressive interactions. Instead, recovering community function required the coordinated replacement of multiple members, and no genomic, taxonomic or pairwise metric predicted which triads would succeed. Community conservatism thus finds its limit at the transition from pairwise to higher-order interactions. To explain this limit, we propose that coexisting strains behave as non-equivalent biological tiles whose interaction edges must align for higher-order function to emerge, an organization we term dynamic *biological tessellation*.

## Results

### Phylogenomic structure strictly dictates pairwise ecological roles

To establish the phylogenomic backbone of our system, we sequenced and assembled 80 Bacillaceae genomes: 79 isolates obtained directly from shallow sediment of the Churince system and one external reference strain (*B. subtilis* PY79). Whole-genome Average Nucleotide Identity (ANI) comparisons resolved 12 distinct clusters, five of which accounted for the majority of the collection: *Subtilis*, *Cereus*, *Cohnii*, *Aquimaris*, and *Bacillus infantis* (Fig. 1a; Supplementary Table 1).

**Fig. 1.**
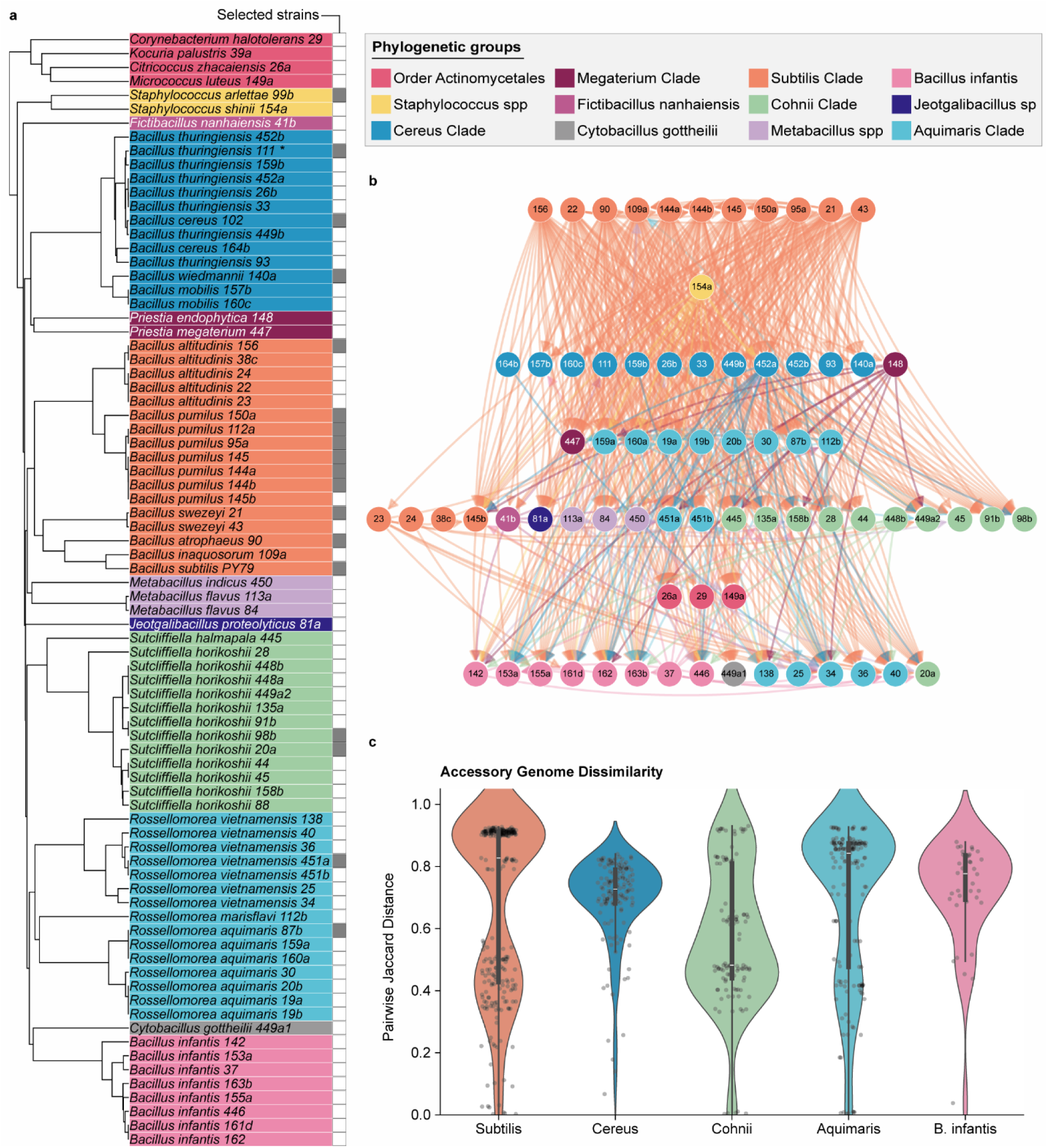
| Phylogenomic conservatism of ecological roles and underlying accessory microdiversity. **a**, Whole-genome ANI dendrogram of the 80 Bacillaceae genomes analyzed. Grey squares denote the 18 representative strains selected for subsequent pairwise and higher-order interaction assays based on phylogenetic relatedness and clade representation. Note: Strain 111 may be identified as either *B. cereus* or *B. thuringiensis* (also in the Cereus clade). We use *B. cereus* 111 (Bc111) for consistency with prior literature. **b**, Eco-genomic interaction network where directed edges represent pairwise antagonistic interactions. Strains are vertically stratified via hierarchical clustering incorporating the Interaction Index (II) and core genomic features (genome size, CDS, GC content, tRNAs, and rRNAs). The vertical axis reflects the community competitive hierarchy, descending from highly antagonistic lineages (top) to susceptible isolates (bottom). Node colors denote phylogenetic clades. Five non-interacting strains were excluded from the network. **c**, Accessory genome dissimilarity within local clades. Pairwise Jaccard distances calculated exclusively on the accessory genome of isolates for the five dominant phylogenetic clades. A distance of 1.0 indicates no overlap in accessory genes. Violins display data distribution; internal boxplots denote medians and interquartile ranges of intra-clade dissimilarity. Individual points represent unique pairwise strain comparisons within each clade. Global pangenome matrices including external reference genomes are provided in Extended Data Fig. 1

To test whether this phylogenetic structure governs interaction potential, we leveraged prior pairwise antagonistic data from solid-medium assays for 78 of these community isolates (Pérez-Gutiérrez *et al.*, 2013). By computing an Interaction Index (II) that integrates antagonistic capacity and susceptibility, we classified the isolates into Antagonistic, Resistant, and Sensitive roles (Supplementary Table 2). We then constructed a directed interaction network spatially stratified by a multivariate clustering of the II alongside core genomic features (Fig. 1b; Supplementary Table 2). Rather than sorting strictly by antagonistic capacity, this multi-trait clustering naturally segregated strains by taxonomic proximity. The antagonistic phenotype was heavily concentrated within the *Subtilis* clade, whereas resistant strains were predominantly associated with the *Cereus* clade, and sensitive isolates were broadly distributed across the *Sutcliffiella*, *Aquimaris*, and *Bacillus infantis* clades. This structural convergence demonstrates that at the pairwise level, ecological capacity and genomic architecture are tightly coupled and conserved along deep phylogenetic boundaries.

### Accessory genome variation reveals contrasting patterns of microdiversity across clades

Having established that ecological roles are conserved across clades, we asked whether strains within these groups are genomically equivalent. We first reconstructed the global pangenomes of the five dominant clades, integrating external reference genomes (Extended Data Fig. 1; Supplementary Table 3). Most reference genomes retained high singleton counts relative to the Cuatrociénegas collection, revealing minimal accessory overlap and underscoring the evolutionary isolation of the local ecosystem. To quantify local microdiversity, we isolated the genomic data of the Cuatrociénegas community, excluded the core genome of each clade, and computed pairwise Jaccard distances exclusively on the accessory gene clusters (Fig. 1c).

If phylogenetic conservatism equated to genomic uniformity, closely related strains sharing an ecological role should exhibit similar accessory profiles. Instead, four of the five dominant clades exhibited high overall accessory dissimilarity, with median pairwise Jaccard distances exceeding 0.7 (indicating less than 30% shared accessory gene content), whereas the *Cohnii* clade showed moderate dissimilarity (median ∼0.5; Fig. 1c). However, the architecture of this variation differed across lineages. While *Bacillus infantis*, *Cereus*, and *Aquimaris* displayed broad, continuous accessory plasticity, the *Subtilis* clade (overall median > 0.8) exhibited a distinct bimodal distribution driven by its internal split into the *B. subtilis* and *B. pumilus* subclades (Fig. 1c; Extended Data Fig. 1a). The upper distribution peak reflects the deep divergence between these two subclades. Conversely, the lower peak corresponds to comparisons within the *B. pumilus* subclade, where accessory genomes are tightly conserved (with 10 of 12 strains harboring between 0 and 7 singletons; Extended Data Fig. 1a). Thus, our collection spans a wide spectrum of microdiversity, from highly plastic clades (*Cereus*, *Aquimaris*, *B. infantis*) to a tightly cohesive sub-lineage (*B. pumilus*), providing an ideal testbed to evaluate how both broad and minimal genomic variation impact ecological interactions.

### BGC repertoires and pairwise antagonism are conserved by clade but vary quantitatively across strains

To understand how this genomic structure shapes the competitive arsenal of the antagonistic *Subtilis* clade, we profiled biosynthetic gene clusters (BGCs) across the collection. Global profiling confirmed that the *Subtilis* and *Cereus* clades harbor the highest overall BGC counts (Extended Data Fig. 2a). Within the *Subtilis* clade, BGC composition strictly mirrored the bimodal accessory genome structure observed in our Jaccard analysis (Fig. 1c, Fig. 2a). While the *B. subtilis* and *B. pumilus* subclades encode distinct BGC repertoires, strains within the *B. pumilus* subclade share a highly conserved biosynthetic profile, differing from one another by only one or two clusters.

**Fig. 2.**
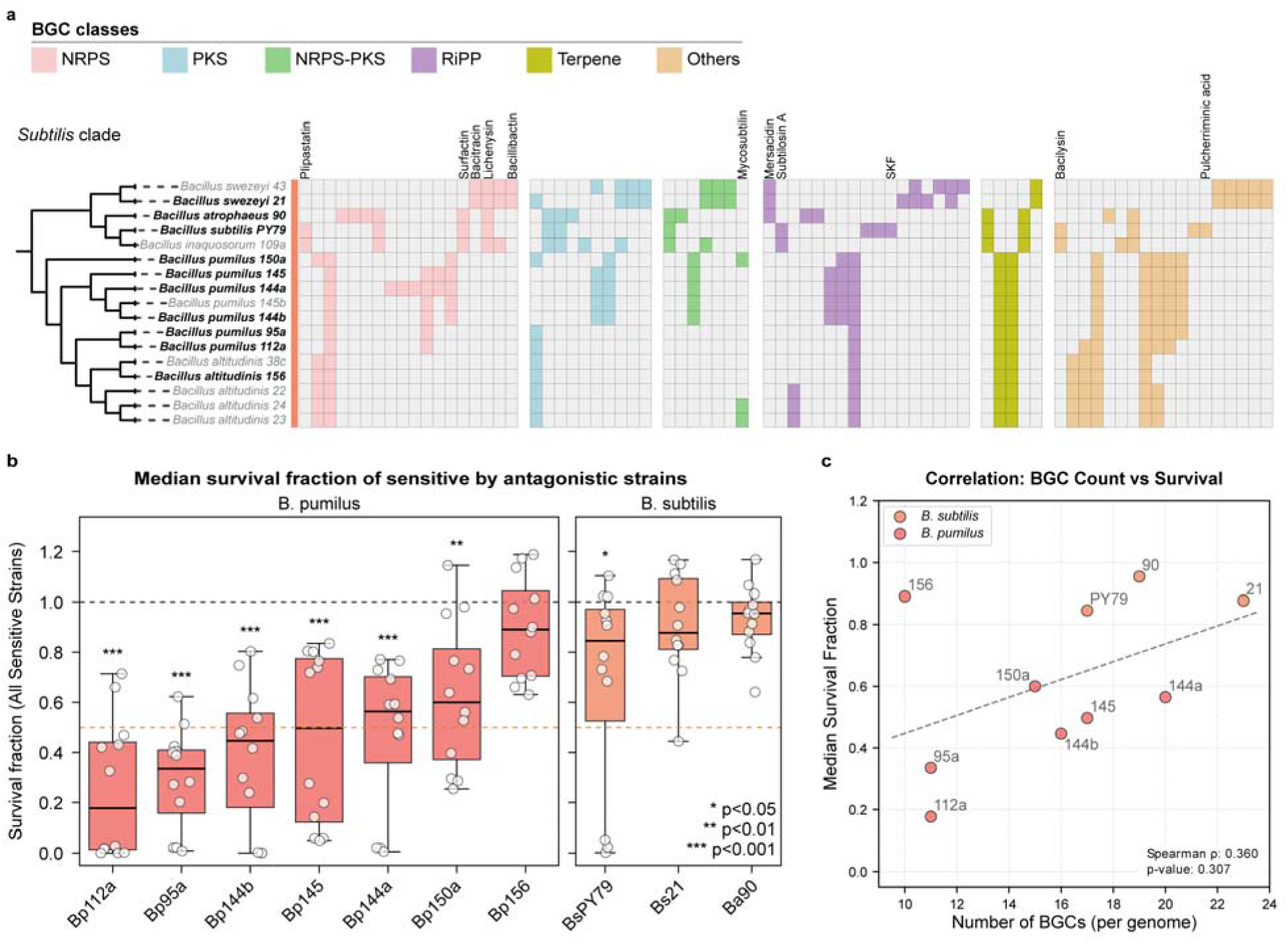
| Biosynthetic microdiversity drives quantitative unpredictability in pairwise antagonism. a, Biosynthetic repertoires within the *Subtilis* clade. Hierarchical clustering based on BGC presence/absence profiles highlights both characterized antimicrobial clusters and uncharacterized BGCs. The distribution reveals highly individualized biosynthetic mosaics even among closely related strains within the *B. pumilus* and *B. subtilis* sub-clades. Strains selected for subsequent interaction assays are indicated in bold. **b,** Quantitative hypervariability of antagonistic strength. Survival fractions (t₃₀/t₀) of sensitive targets following 30 min of liquid co-culture with individual antagonists, grouped by sub-clade. Points represent single observations; boxplots show medians and interquartile ranges. While sub-clade conservatism is evident, individual strain lethality varies drastically. Significance versus a survival fraction of 1.0 (dashed line) was calculated using two-sided Wilcoxon signed-rank tests with Benjamini–Hochberg correction; asterisks denote adjusted *P* values (*q*). **c,** Decoupling of absolute biosynthetic potential and ecological function. The median survival fraction of sensitive targets (y-axis) plotted against the total BGC count per genome (x-axis) for each antagonist. Points are colored by sub-clade; the dashed line indicates the linear trend. The lack of a significant relationship (Spearman’s ρ = 0.36, *P* = 0.307) demonstrates that antagonistic capacity is not an additive trait driven by total BGC burden.

We next tested whether strains with such conserved BGC profiles behave equivalently during direct competition. We characterized pairwise interactions between ten Subtilis-clade strains against four sensitive strains in well-mixed liquid co-cultures, an environment that recapitulated the qualitative outcomes of solid-medium assays (Extended Data Fig. 2b–f). Qualitatively, phylogenetic conservatism held across all ten *Subtilis*-clade antagonists, as every strain inhibited the sensitive targets. Quantitatively, however, survival fractions revealed striking strain-specific variation (Fig. 2b; Supplementary Table 4). Although sub-clade conservatism was evident, with *B. pumilus* strains consistently producing stronger inhibition than *B. subtilis* strains (Mann–Whitney U test, P = 6.98 × 10⁻□), individual *B. pumilus* isolates exhibited markedly different killing strengths across specific sensitive targets despite sharing nearly identical BGC profiles and accessory genomes (Fig. 2b; Extended Data Fig. 2f).

Furthermore, total BGC count per genome did not correlate with antagonistic strength across the Subtilis clade (Spearman’s *p* = 0.36, P = 0.307; Fig. 2c). Together, these results demonstrate that competitive dominance is not an additive trait dictated by total BGC number. Even within the genomically cohesive B. pumilus subclade, subtle variation in BGC composition or regulatory deployment drives substantial quantitative differences in pairwise antagonism, raising the question of whether these fine-scale differences completely disrupt functional interchangeability in higher-order communities.

### Phylogenetic conservatism breaks down at the transition to higher-order interactions

To test whether strain-level variation disrupts functional interchangeability in complex communities, we used the synthetic three-species BARS model to quantify the emergent protection of a sensitive strain by a resistant community member against a lethal antagonist (Fig. 3a). To ensure a sufficiently strong antagonistic baseline for detecting emergent rescue, we focused our higher-order assays on the four *B. pumilus* isolates (Bp145, Bp95a, Bp112a, and Bp144b) that reduced median target survival below 0.5 in liquid co-cultures (Fig. 2b; Supplementary Table 5). We also confirmed that the four candidate resistant strains lacked independent antagonistic effects against the sensitive targets across pairwise assays, establishing a neutral baseline (Extended Data Fig. 3a; Supplementary Table 6).

**Fig. 3.**
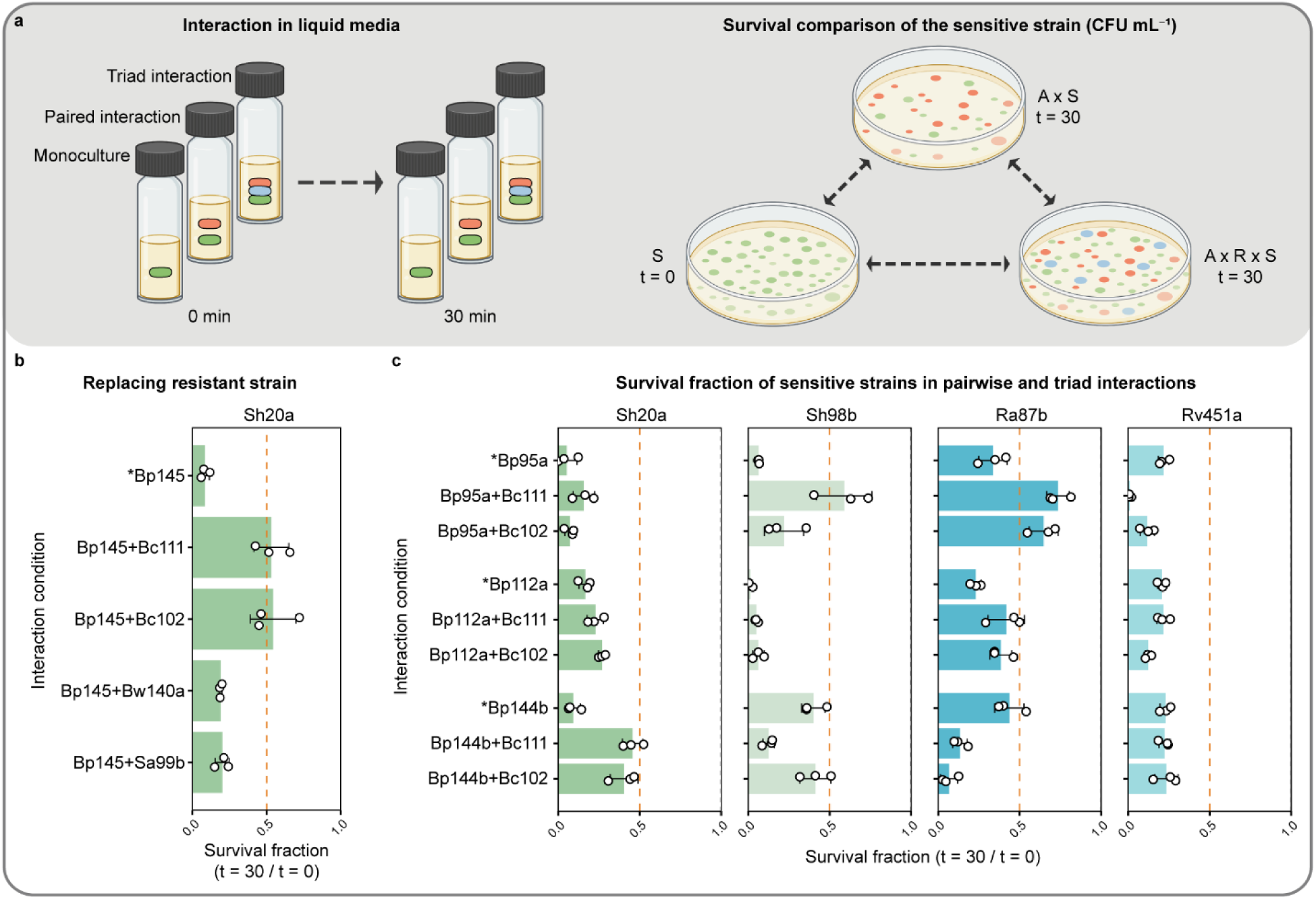
| Higher-order protective interactions are highly context-dependent. **a,** Workflow of the liquid co-culture assay. Sensitive strain survival was quantified by comparing colony-forming units (CFU mL⁻¹) in monoculture at 0 min with CFUs after 30 min of co-culture in either pairwise (antagonist) or higher-order (antagonist + resistant) configurations. **b,** Survival of Sh20a when alternative resistant strains were substituted into the canonical BARS triad (Sh20a × Bp145 × Bc111). **c,** Survival of sensitive strains (Sh20a, Sh98b, Ra87b, and Rv451a) co-cultured with antagonists (Bp95a, Bp112a, and Bp144b), alongside corresponding triads including a resistant strain (Bc111 or Bc102). The black dashed line marks the monoculture baseline (value = 1); the red dashed line indicates the 0.5 survival threshold. Bars represent mean ± s.d. (n = 3 independent replicates). Significance was assessed via one-or two-sample t-tests (Supplementary Table 7) Asterisk denote pairwise interaction. Raw CFU ml^-1^ counts are available via Zenodo (DOI: 10.5281/zenodo.23126602).

We then tested the core assumption of community conservatism, whether strains sharing the same pairwise ecological role and clade membership are functionally interchangeable. We systematically substituted the resistant member of the canonical BARS triad (Bp145 × Sh20a × Bc111) with alternative resistant isolates, predominantly from the highly plastic *Cereus* clade (Bc102 and Bw140a) alongside Sa99b (Fig. 3b). Strikingly, functional redundancy was the exception rather than the rule; among the candidates, only *B. cereus* 102 (Bc102) restored protection against Bp145 to levels comparable to the original *B. cereus* 111 (hereafter Bc111) strain (Fig. 3b).

More critically, the ability of Bc102 to substitute Bc111 was strictly context-dependent and failed to generalize across alternative antagonistic threats (Fig. 3c). While both resistant strains successfully protected Sh20a from the antagonist Bp144b, they failed completely to rescue it from Bp95a or Bp112a. Similarly, Bc111 protected Sh98b from Bp95a whereas Bc102 did not, and neither resistant strain rescued Rv451a under any tested condition (Fig. 3c; Extended Data Fig. 3b). This asymmetry exposes the limit of taxonomic and genomic prediction: Bp145 and Bp95a belong to the same cohesive subclade, share nearly identical accessory genomes and BGC repertoires (Fig. 1c, Fig. 2a), and produce severe killing in pairwise assays (Fig. 2b); yet, they elicit entirely opposite higher-order outcomes when confronted with the exact same resistant partner. This demonstrates that protective equivalence is not an intrinsic, conserved trait of a clade, but an emergent property unique to specific, highly constrained strain combinations.

### Community outcomes are non-additive emergent properties restricted by biological tessellation

To determine whether these context-dependent outcomes represented true higher-order effects rather than additive pairwise interactions, we implemented a quantitative additive null model (Fig. 4a; Supplementary Table 7). In this framework, expected triad survival equals the baseline pairwise antagonistic effect, assuming no direct effect from the resistant strain. Strikingly, across all 34 triads evaluated, community outcomes frequently and severely deviated from pairwise expectations (Fig. 4a). Thirteen combinations (38%) displayed significant emergent rescue, with positive deviations reaching up to an order of magnitude above null predictions. Conversely, five combinations exhibited suppressive higher-order interactions, where triad survival fell significantly below the pairwise baseline. A multiplicative null model yielded the identical qualitative classification across all 34 triads, confirming that these dynamics are true emergent phenomena and not artifacts of the additive assumption (Supplementary Tables 8, 9).

**Fig. 4.**
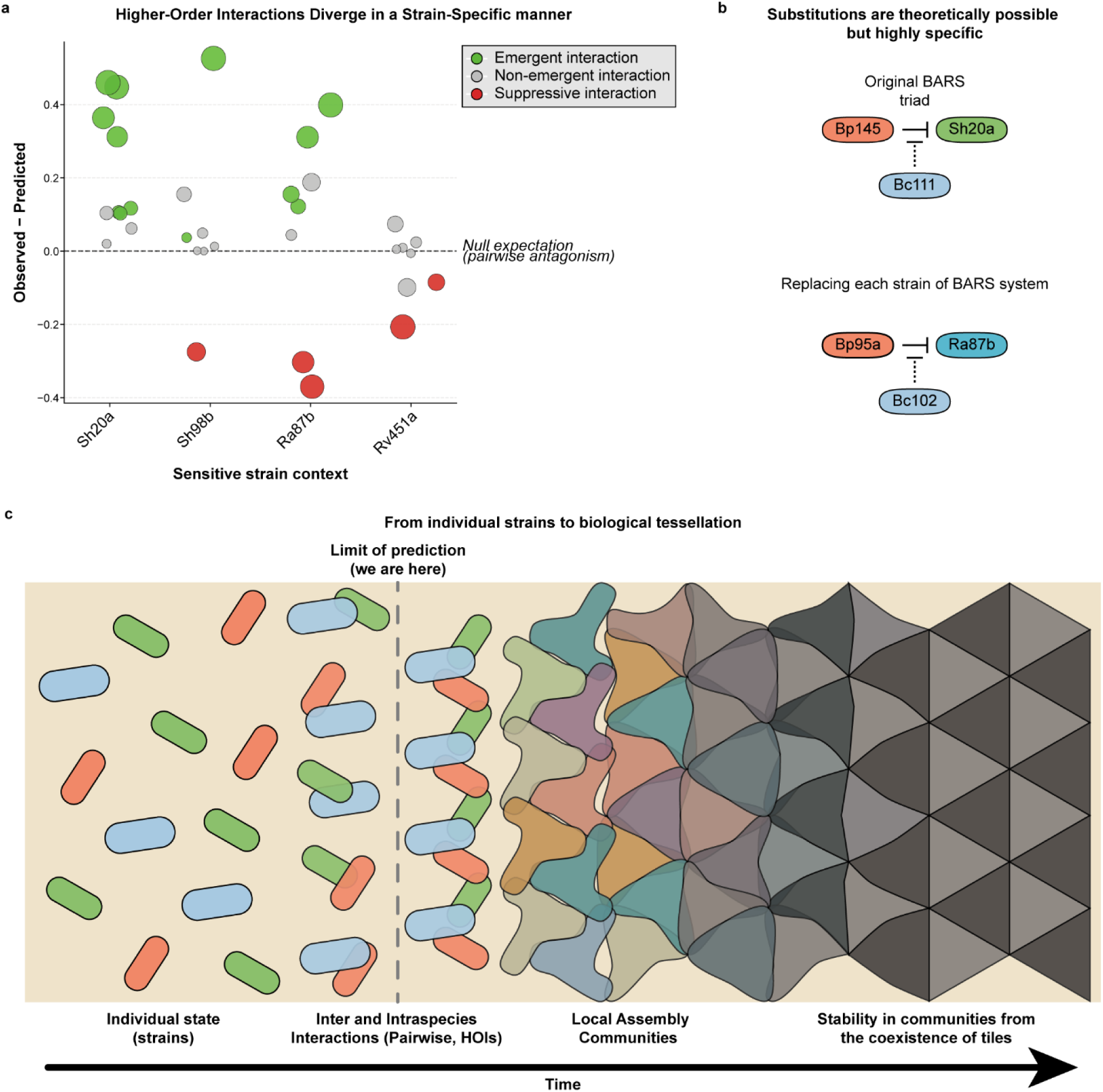
| Community outcomes are non-additive emergent properties consistent with biological tessellation and its hypothesized dynamics. a Breakdown of pairwise additivity in higher-order interactions. Divergence between observed triad survival and additive null model predictions for all 34 three-member communities. Points represent unique strain combinations, ordered by sensitive strain context (Observed − Predicted). The horizontal dashed line denotes expected survival under purely additive pairwise antagonism. Positive deviations (green) indicate significant emergent rescue, whereas negative deviations (red) denote suppressive higher-order interactions. Gray points represent non-significant outcomes. Point size scales with the absolute magnitude of deviation. **b,** Recovery of community function through systemic reconfiguration. Schematic of an alternative functional BARS triad (Bp95a × Ra87b × Bc102) that successfully achieves emergent protection despite the complete substitution of all three original community members, showing that functional redundancy can be recovered through the joint reconfiguration of all members, even when isolated single-strain swaps fail. **c,** Proposed conceptual model of dynamic biological tessellation. While phylogenetic conservatism reliably predicts individual traits and pairwise outcomes, the transition to higher-order interactions (dashed vertical line) marks a sharp limit of prediction. Each strain is depicted as a non-equivalent tile; beyond this threshold, only strains whose interaction edges align assemble into functional higher-order modules. Local communities and broader ecosystem stability are hypothesized to emerge from the interlocking and dynamic coexistence of these compatible tiles over time, rather than from the free substitution of individual strains.

The failure of single-strain substitutions (Fig. 3) coupled with pervasive non-additivity (Fig. 4a) suggests that functional interchangeability requires the synergistic alignment of multiple genomes. Consistent with this, protection was recovered in an entirely alternative strain combination (Bp95a × Ra87b × Bc102), in which all three original members were replaced by strains from their respective clades (Fig. 4b). Although this represents a single combination, it provides an existence proof that role-based functional equivalence can be achieved and suggests that it may depend on the joint configuration of the triad rather than on swapping individual parts into a pre-existing module.

Despite being drawn from clades with conserved ecological roles, most tested strain combinations failed to reconstitute the protective phenotype, exposing a sharp predictability boundary at the transition from pairwise to higher-order interactions. To conceptualize how communities assemble beyond this threshold, we propose a framework of dynamic biological tessellation (Fig. 4c). In this model, individual strains do not integrate into communities as freely interchangeable units; instead, each strain behaves as a non-equivalent tile whose genomic and physiological configuration defines its interaction edges, and higher-order function emerges only from combinations whose edges align. We hypothesize that local community assembly and ecosystem stability emerge from the spatial and metabolic interlocking of these compatible tiles rather than from a random assortment of redundant species. Because strains reshape their own interaction edges as they modify their environment, and because individual strains turn over, community stability beyond this higher-order threshold may shift dynamically; predicting emergent microbiome properties require identifying the specific multi-strain tessellations that govern assembly.

## Discussion

Predicting ecological functions from genomic data remains a central goal in microbial ecology, yet our results expose a fundamental, scale-dependent limit to this objective. Phylogenetic trait conservatism provides a robust baseline: ecological roles and BGC repertoires are deeply organized by lineage in the Cuatrociénegas *Bacillaceae* (Pérez-Gutiérrez *et al.*, 2013; Gómez-Lunar *et al.*, 2016; Rodríguez-Torres *et al.*, 2017; González-Sánchez *et al.*, 2026). At the pairwise scale, this conservatism reliably predicts interaction potential and niche boundaries (Losos, 2008; Wiens *et al.*, 2010). However, this predictive power collapses precisely where community function ultimately emerges in higher-order interactions. Crucially, this breakdown occurs across the entire spectrum of intraspecific variation—whether substituting resistant strains from clades with high accessory genome plasticity (Cereus) or swapping antagonists from a cohesive sub-lineage sharing nearly identical accessory genomes and BGC repertoires (*B. pumilus*). When strains equivalent by core phylogeny, biosynthetic potential, and pairwise ecological role are assembled into a three-species context, their realized ecological functions diverge sharply. This demonstrates that strain-level variation does not merely introduce background noise; it defines the rules translating genomic potential into realized ecology, and these rules change strictly with the scale of interaction.

The significance of this predictability barrier is amplified by the pervasive nature of conservatism in our system. In the El Churince *Bacillaceae*, conservatism is not a weak statistical trend but a deep organizational constraint governing antagonistic networks, stress responses, and biosynthetic repertoires. Even community-wide responses to severe environmental desiccation in this system rely on ancestrally shared physiological programs (González-Sánchez *et al.*, 2026). This makes the failure of genomic prediction at the higher-order interaction scale particularly revealing. Our findings do not contradict niche conservatism but reframe it as scale-dependent: conservatism governs the broad ecological theater and the baseline capacity to interact, while strain-level microdiversity determines the specific performance that secures local stability. This aligns with the principle that functional redundancy decreases as functional complexity increases (Jia & Whalen, 2020). Higher-order interactions, contingent on the simultaneous interplay of three species rather than a single isolated trait, represent exactly such a complex function.

The mechanistic basis of this predictive breakdown lies in the integrated genomic context required to deploy competitive traits. Closely related *B. pumilus* strains encoding nearly identical BGC repertoires display markedly different killing strengths in pairs and elicit opposite outcomes in triads, implying that ecological function depends not merely on gene presence, but on the regulatory and metabolic configuration of the whole genome (Casadesús & Low, 2006; González-Torres *et al.*, 2015). While complex traits like antagonistic BGC machinery are expected to be deeply conserved (Martiny *et al.*, 2015), our results show they vary functionally at the microevolutionary scale. The BGCs present across a clade dictate what is metabolically possible for the lineage, but each strain deploys its chemical arsenal differently depending on its regulatory wiring and the specific cues of its interacting partners. Indeed, our null model analysis shows that introducing a third, pairwise-neutral strain can either attenuate antagonism to produce emergent rescue or, conversely, exacerbate killing in suppressive higher-order interactions. Consequently, genes do not act as isolated units. When scaled to the community level, ecosystem performance becomes a non-additive, emergent property of the specific interaction network (Kamrad *et al.*, 2026; Moraïs *et al.*, 2026), explaining why genomic data can identify broad functional categories but cannot specify which strain combinations will yield a viable community outcome (Widder *et al.*, 2016; Goyal *et al.*, 2022).

Natural communities comprise multiple overlapping interaction modules. If each module is subject to the stringent compatibility constraints observed in our BARS model, where single-strain swaps consistently failed and fewer than half of the tested combinations produced emergent rescue, the space of successful community configurations is combinatorially restricted (Friedman *et al.*, 2017; Sanchez-Gorostiaga *et al.*, 2019; Gallardo-Navarro *et al.*, 2025). Notably, the recovery of protection in a completely reconfigured triad (Bp95a × Ra87b × Bc102) shows that functional equivalence is attainable through a new strain combination, even when single substitutions fail. Together with the failure of single-strain swaps, this observation leads us to hypothesize that functional equivalence resides at the level of the multi-strain module rather than the individual organism, a prediction that can be tested by assembling a larger set of fully reconfigured triads. We conceptualize this organization as a dynamic biological tessellation, in which strains behave as non-equivalent tiles. Each tile’s genomic and physiological configuration defines its active interaction edges, and higher-order function emerges only when these edges align across coexisting members; the community’s realized structure is the mosaic of such compatible modules.

The BARS assay captures the formation of these modules within a 30-minute window before cell division and broader metabolic remodeling occur. The fact that phylogenetic conservatism fails to predict higher-order outcomes even within this brief window indicates that strict strain-level compatibility governs assembly from the very first contact. Beyond this window, tiles reshape their own edges as they modify their chemical and physical surroundings, as illustrated by the dynamically induced spatial segregation of motile isolates from this system (Gallardo-Navarro *et al.*, 2025). We therefore expect tessellations to be transient configurations that assemble and disassemble as strains change their phenotypes and environment, making microbiome stability even harder to predict from species-level inventories.

This framework carries a cautionary implication for the current era of large-scale metagenomics. The capacity to sequence and taxonomically classify thousands of strains creates a powerful, yet potentially misleading, sense of ecological understanding. Knowing that members of a species encode specific biosynthetic capabilities is insufficient to predict their behavior in a community context. For instance, the rational design of microbial consortia for agriculture or biocontrol typically relies on selecting strains by species identity and individual functional profiles. Our findings indicate that such approaches will inevitably fail to predict which specific strains will produce the desired emergent community outcome; experimental validation of precise strain combinations remains indispensable. More broadly, just as the loss of ecological interactions can precede and outpace species loss (Valiente-Banuet *et al.*, 2015), a mismatch of interaction edges between conspecific strains can abolish community function without any change in species composition. Predictive frameworks will therefore need to shift from a gene-centric to an interaction-edge view of community assembly.

Ultimately, microdiversity must be recognized not as background noise, but as a fundamental determinant of ecological dynamics. Recognizing this shifts the focus of microbial ecology from identifying *who is there* to understanding *who interacts with whom*, and under what specific conditions. Phylogenetic conservatism tells us which tiles are available; only the alignment of their interaction edges determines which communities can assemble.

## Methods

### Strain selection

Eighteen bacterial strains were selected from a culture collection of isolates from the Cuatrociénegas Basin, Coahuila, Mexico, previously characterized for pairwise ecological interactions (Pérez-Gutiérrez *et al.*, 2013). Strains were chosen to represent the three ecological roles defined in the BARS model (Aguilar-Salinas & Olmedo-Álvarez, 2023): antagonistic (A, n = 10), resistant (R, n = 4), and sensitive (S, n = 4). Full strain metadata are provided in the Supplementary Table 1.

### Spot-on-lawn antagonism assays

Antagonistic interactions were assessed on solid marine medium using a spot-on-lawn design in which each strain was tested both as a lawn and as a spotted isolate against all other strains. Lawns were prepared by spreading 100 µl of overnight culture (∼1 × 10 CFU ml⁻¹) onto square plates (15 × 15 cm), and 12 µl of each overnight culture was spotted onto every lawn. Plates were incubated at 28°C for 24–48 h, depending on halo visibility. Inhibition zones were measured as the distance (mm) between the edge of the spotted colony and the edge of the visible clearing halo. All assays were performed in four independent replicates.

### Pairwise interaction assays in liquid medium

Interaction assays were performed in liquid marine medium (MM-L). Overnight cultures were grown for 14 h at 28°C with agitation at 180 rpm and subcultured into fresh medium until reaching exponential phase (∼10 CFU ml⁻¹ for S strains and ∼10 CFU ml⁻¹ for A and R strains), monitored by absorbance on a Klett-Summerson colorimeter (red filter). All co-cultures were assembled in glass vials to a final volume of 10 ml as follows: S×A (6 ml S, 2 ml A, 2 ml MM-L), S×R (6 ml S, 1 ml R, 3 ml MM-L), and A×R (2 ml A, 1 ml R, 7 ml MM-L). Vials were mixed gently by hand every 10 min throughout the 30 min interaction period. CFU ml⁻¹ were determined by serial dilution in PBS and plating on solid marine medium (MM-S) at t₀ and t₃₀, followed by incubation at 28°C for 48 h. Strains were distinguishable by colony morphology. Colonies were counted using ImageJ. Survival fractions of sensitive strains were calculated as CFU ml⁻¹ at t₃₀ in co-culture relative to their monoculture at t₀ for each biological replicate. All interactions were performed in at least three independent replicates. Survival fractions are provided in Supplementary Tables 4 and 6.

### Three-member interaction assays

Three-member community assays followed the same protocol as the pairwise interaction assays, with co-cultures assembled as 6 ml S, 2 ml A, and 1 ml R, with 1 ml MM-L to a final volume of 10 ml. Survival fractions of sensitive strains were calculated as described above. Statistical survival fractions are provided in Supplementary Table 7 as “Observed S_survival”. Raw CFU ml⁻¹ counts of every paired and triad interaction are available via Zenodo (DOI: 10.5281/zenodo.23126602).

### DNA extraction and whole-genome sequencing

Genomic DNA was extracted from 51 bacterial isolates using a commercial kit following the manufacturer’s protocol. The remaining 29 isolates had been previously sequenced and genome assemblies are publicly available under the accession numbers listed in Supplementary Table 1. Whole-genome sequencing was performed on an Illumina MiSeq and NovaSeq platform using paired-end libraries prepared according to the manufacturer’s guidelines. Sequencing coverage was estimated using the Lander–Waterman equation. Raw reads were quality-assessed with FastQC v0.12.1 and trimmed with Trimmomatic v0.39; full trimming parameters are described in Supplementary Materials and Methods.

### Genome assembly and functional annotation

Quality-filtered reads were assembled de novo using SPAdes v3.13.0 (Bankevich *et al.*, 2012) under the De Bruijn graph framework, using k-mer sizes of 21, 33, 55, 77, 99, and 127. Assembly statistics, including contig length distribution and total assembly size, were subsequently calculated for each genome assembly.

Structural and functional genome annotation was performed using Prokka v1.14.6 (Seemann, 2014). Functional assignments of predicted coding sequences (CDSs) were based on hierarchical searches against curated protein databases, including UniProtKB/SwissProt, ISfinder, and the NCBI Bacterial Antimicrobial Resistance Reference Gene Database. This approach enabled the annotation of conserved housekeeping genes, mobile genetic elements, and putative antimicrobial resistance determinants across the genome dataset. For taxonomy classification see Supplementary Information. All data were submitted to NCBI database (Supplementary Table 1).

### Average Nucleotide Identity (ANI) analysis

Pairwise genomic similarity among the analyzed 80 genomes was assessed through an all-versus-all average nucleotide identity (ANI) analysis. ANI values were calculated using a BLAST-based approach (ANIp) implemented in PERL (calcANI.pl at https://github.com/Computational-conSequences/SequenceTools), generating a pairwise identity matrix that was subsequently used for comparative genomic analyses. The raw all-versus-all comparisons are available via Zenodo (DOI: 10.5281/zenodo.23126602).

### Pangenomic characterization and comparative analysis

To establish a robust comparative framework, reference genomes were retrieved and integrated with the 80 previously sequenced bacterial isolates, bringing the total dataset to 109 genomes. Prior to database generation, reference FASTA sequences were reformatted and filtered to remove contigs shorter than 1,000 bp using *anvi-script-reformat-fasta* to ensure assembly quality. For each genome, a contigs database was generated using *anvi-gen-contigs-database* from the Anvi’o v8 software suite (Eren *et al.*, 2020). Subsequently, all databases were annotated for single-copy core genes using *anvi-run-hmms*. The 109 genomic databases were then unified into a single centralized genomes storage file (*anvi-gen-genomes-storage*).

The global pangenome was computed utilizing the *anvi-pan-genome* workflow. During this step, open reading frames were translated, and amino acid sequence similarities were calculated across all genomes using the default DIAMOND search. Homologous sequences were then clustered into discrete gene families (gene clusters) using the Markov Cluster Algorithm (MCL) with a default inflation parameter of 2.0. To assess genomic similarity and establish phylogenetic relationships among the strains, hierarchical clustering of the 109 genomes was explicitly enforced *(--enforce-hierarchical-clustering*) based on the frequencies of shared gene clusters, employing Euclidean distance and Ward’s linkage method.

For downstream differential comparisons and targeted intersection visualization, the global presence-absence matrix of the 109 genomes was subsetted to focus exclusively on the five most highly represented lineages (*Subtilis* Clade, *Cereus* Clade, *Bacillus infantis*, *Aquimaris* Clade, and *Cohnii* Clade). This filtering step highlighted major differential genomic architectures while retaining the biological context of the global clustering structure.

### Accessory genome dissimilarity and microdiversity analysis

To quantitatively assess strain-level microdiversity, we calculated pairwise dissimilarities exclusively among the local Cuatrociénegas isolates. The global pangenome presence-absence matrix generated by Anvi’o was first filtered to remove all external reference genomes. For each of the five dominant phylogenetic clades (*Subtilis*, *Cereus*, *Cohnii*, *Aquimaris*, and *Bacillus infantis*), the local core genome—defined as gene clusters present in 100% of the focal clade’s environmental isolates—was excluded. The remaining clade-specific accessory gene clusters were used to compute a pairwise Jaccard distance matrix. This distance (where 0 indicates identical accessory repertoires and 1.0 indicates no shared accessory gene clusters) was calculated across all unique strain pairs within each clade to evaluate intra-clade dissimilarity distributions

### Biosynthetic gene cluster profiling

Biosynthetic gene clusters (BGCs) were predicted across the 80 high-quality genomes using antiSMASH v7.0 (Blin *et al.*, 2023). To evaluate biosynthetic similarity among genomes, predicted BGCs were subsequently clustered into Gene Cluster Families (GCFs) and Gene Cluster Clans (GCCs) using BiG-SCAPE v1.1.9 (Navarro-Muñoz *et al.*, 2020). BiG-SCAPE calculates pairwise distances among BGCs based on a weighted combination of the Jaccard Index (JI), adjacency index (AI), and domain sequence similarity (DSS), enabling the identification of related biosynthetic architectures across genomes.

GCFs and GCCs were defined using distance cutoffs of 0.3 and 0.7, respectively. The resulting presence–absence matrix of biosynthetic cluster groups across genomes was further analyzed using hierarchical clustering with average linkage methods and linear modeling approaches to assess patterns of biosynthetic diversity and genomic relatedness among strains (For complete matrix see Source Data). To see complete BGCs count distributions across 80 strains see Supplementary Figure 3.

### Interaction Index and network analysis

Pairwise antagonistic interactions were experimentally quantified using spot-on-lawn assays, yielding a binary directed adjacency matrix representing 6,006 interaction outcomes across 78 focal strains (matrix available via Zenodo [DOI: 10.5281/zenodo.23126602]). To evaluate competitive boundaries, we derived a Competition Index (*CI = AI - SI*), where the Antagonism Index (*AI*) and Sensitivity Index (*SI*) represent the proportion of the community a focal strain inhibits (out-degree) or is inhibited by (in-degree), respectively. This metric was subsequently normalized against the maximum observed antagonistic potential and linearly rescaled to a [-1,1] interval to establish the final Interaction Index (*II*). Strains were categorically assigned to antagonistic (*II > 0.4*), resistant (-*0.4 ≤ II ≤ 0.4*), or sensitive (II < -0.4) roles based on empirical discontinuities along the rescaled gradient.

To spatially stratify the community, a directed interaction network was rendered utilizing NetworkX and Matplotlib. Network topology was strictly governed by a multivariate hierarchical clustering of the II alongside core genomic features (genome size, CDS, GC content, rRNAs, and tRNAs). Eco-genomic features were standardized via Z-score normalization using scikit-learn (StandardScaler), followed by the computation of Euclidean distances and Ward’s minimum variance clustering implemented in SciPy. The optimal taxonomic stratification was definitively resolved by maximizing the Silhouette score.

### Additive and multiplicative null models of interaction outcomes

To evaluate whether higher-order interaction outcomes could be predicted from pairwise effects, we compared observed survival fractions with predictions from additive and multiplicative null models. Under the additive model, the combined effect of multiple strains was defined as the sum of their individual pairwise survival fractions relative to monoculture. Under the multiplicative model, combined effects were defined as the product of pairwise survival fractions, assuming independent and non-interacting contributions. Deviations between observed and predicted survival fractions were used to quantify emergent higher-order effects. Statistical evaluation of these deviations is described in the Statistical analysis section. Full model derivations are provided in Supplementary Materials and Methods.

### Statistical analysis

Survival fractions of sensitive strains were compared against a reference value of 1 using one-sample t-tests for each pairwise interaction (n = 3 biological replicates). For comparisons of survival fraction distributions between antagonist clades, a two-sided Mann–Whitney U test was used. For within-strain comparisons against a reference value of 1 across individual antagonistic strains, two-sided Wilcoxon signed-rank tests were applied. Emergence in three-member interactions was assessed by comparing observed survival fractions against additive and multiplicative null model predictions using two-sided one-sample t-tests, and by comparing triad versus pairwise survival fractions using two-sample t-tests. All multiple comparisons were corrected using the Benjamini–Hochberg procedure; adjusted P values (q) < 0.05 were considered statistically significant. Correlation analyses were performed using Spearman’s rank correlation coefficient. All experiments were performed with at least three independent biological replicates. Data are presented as mean ± s.d. unless otherwise indicated in the figure legends.

## Data Availability

Strains isolates are available upon request. All data are available in Supplementary Information, Supplementary Tables, Source Data and via Zenodo (DOI: 10.5281/zenodo.23126602).

## Acknowledgements

We thank África Islas Robles, Valeria Guzmán, Bernardo Aguilar, Eliane Piñón, Roberto Delgado, and Bladimir Villegas for technical work and members of the Olmedo-Alvarez group for helpful discussions. We thank José Antonio Cisneros for his help with photographs.

## Author Contributions

Conceptualization: E.C.M. and G.O.A. Methodology: E.C.M., M.N.M., and G.M.H. Software: E.C.M., M.N.M., V.M.H.A., E.D.B.D and M.H.R. Validation: E.C.M., M.N.M., J.R. and G.O.A. Formal Analysis: E.C.M. and O.R.S. Investigation: E.C.M. and M.N.M. Resources: G.O.A. (laboratory support), V.M.H.A. and E.D.B.D (sequencing platform) Data Curation: E.C.M. and M.N.M. Original Draft: E.C.M. and G.O.A. Review & Editing: E.C.M., J.R., M.H.R. and G.O.A. Visualization: E.C.M., and M.N.M. Supervision: M.H.R. and G.O.A. Project Administration: E.C.M. and G.O.A. Funding Acquisition: M.H.R. and G.O.A.

## Extended Figures

**Extended Data Fig. 1.**
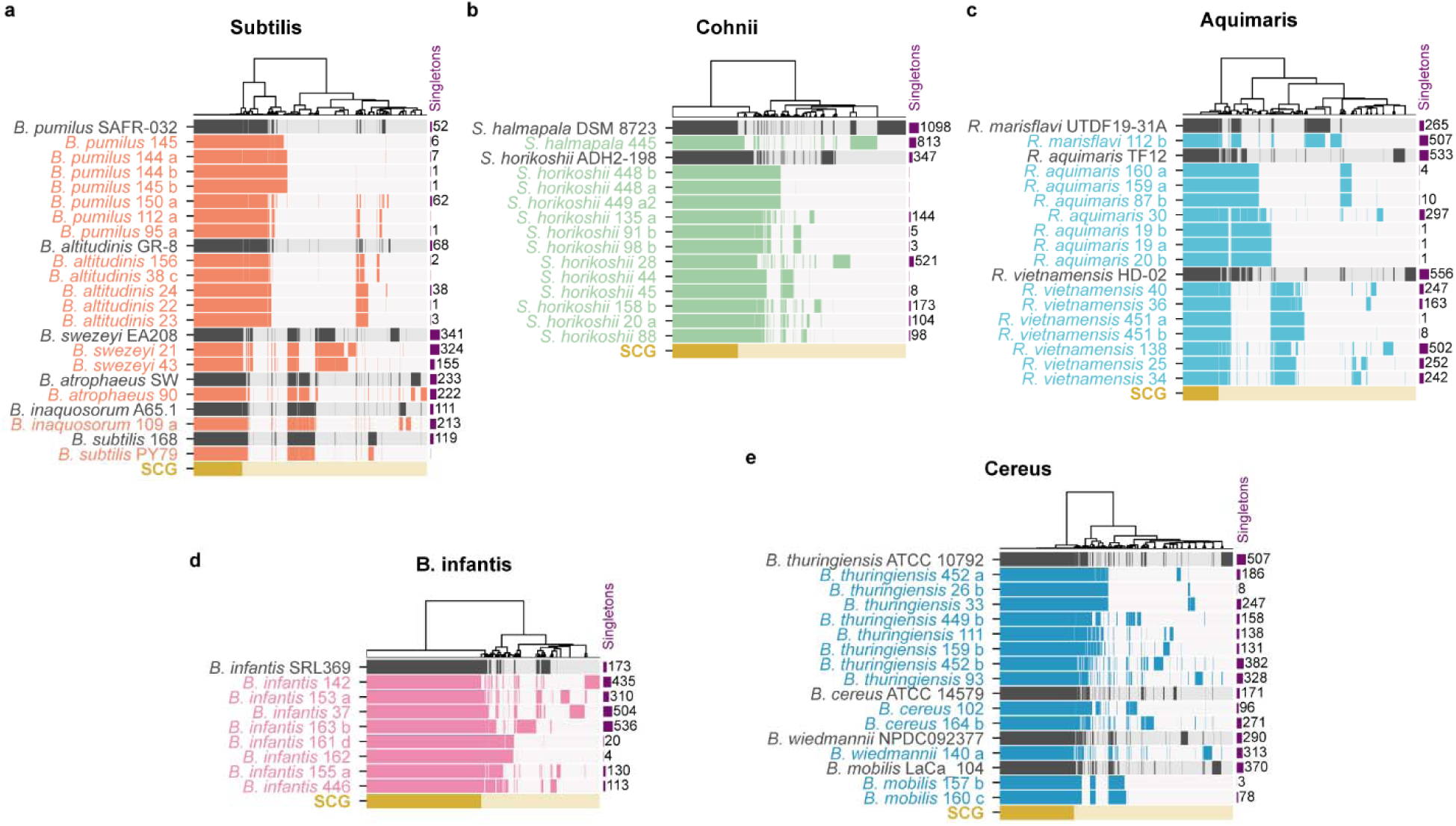
| Pangenomic architecture across dominant *Bacillus* clades. Pangenome architectures of the five dominant clades: Subtilis (**a**), Cohnii (**b**), Aquimaris (**c**), *Bacillus infantis* (**d**), and Cereus (**e**), reconstructed using Anvi’o. Each panel displays a gene-cluster presence/absence matrix, where rows represent individual genomes (reference genomes highlighted in grey) and columns represent hierarchically clustered gene clusters. The bottom track (yellow) indicates single-copy core genes, whereas purple columns denote strain-specific gene clusters. See source data for complete Pangenome matrix.

**Extended Data Fig. 2.**
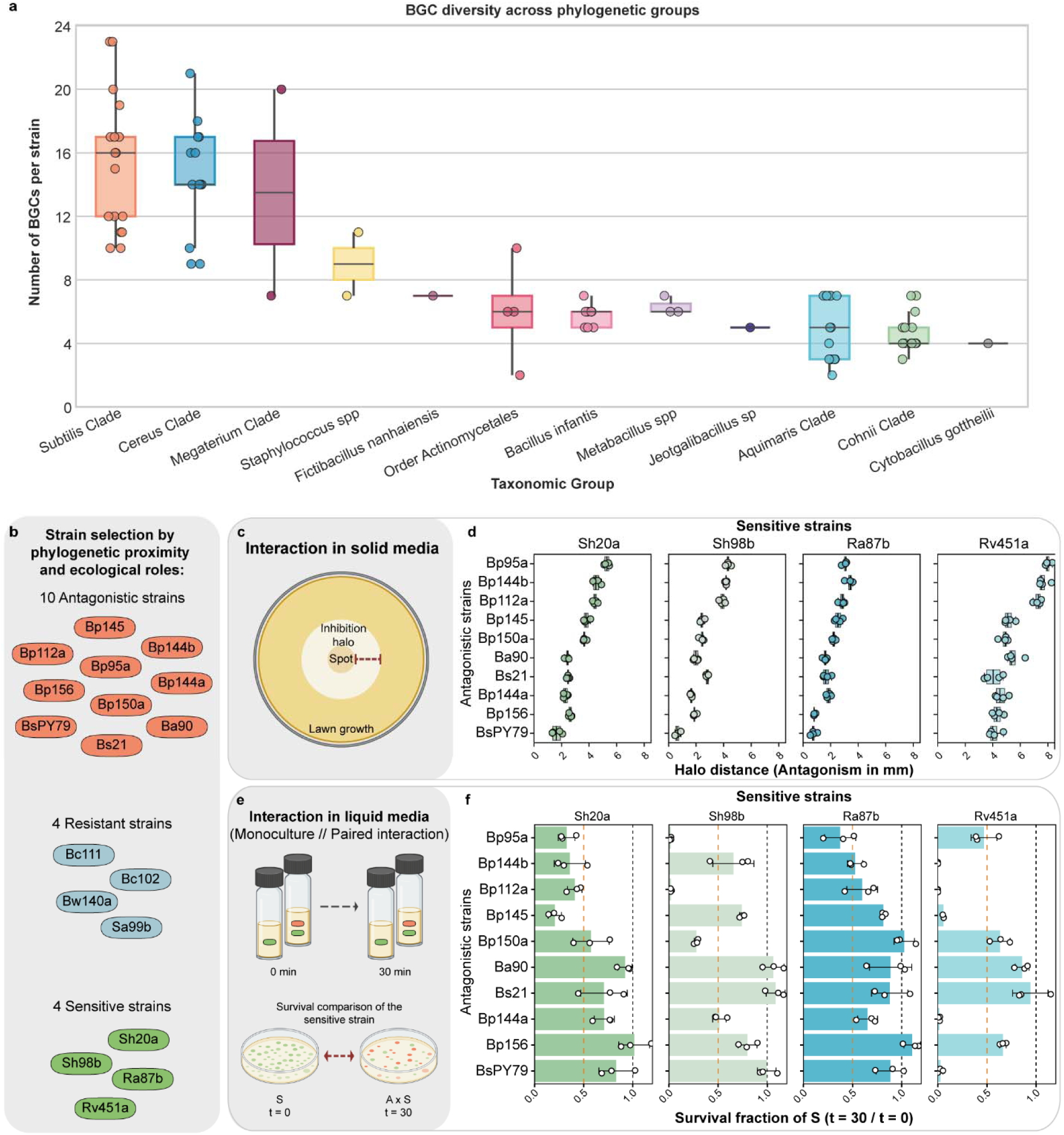
| Global biosynthetic potential and qualitative conservation of pairwise antagonism across spatial contexts. **a**, Biosynthetic gene cluster (BGC) abundance across phylogenetic groups. Points represent individual strains; boxplots denote medians and interquartile ranges, ordered by median BGC count. **b**, The 18-strain subset selected for experimental assays based on phylogenetic relatedness and eco-genomic roles, comprising ten antagonistic, four resistant, and four sensitive strains (details in Supplementary Information). **c**, Schematic of the structured spatial assay (spot-on-lawn). **d**, Inhibition halo diameters (mm) produced by *Subtilis*-clade antagonists against four sensitive targets (Sh20a, Sh98b, Ra87b, and Rv451a). Points represent independent biological replicates; boxplots show medians and interquartile ranges. **e**, Schematic of the well-mixed liquid co-culture assay. Survival fractions represent the ratio of sensitive colony-forming units (CFU mL⁻¹) at 30 min relative to the t₀ monoculture baseline. **f**, Survival fractions in liquid co-culture perfectly recapitulate the qualitative outcomes of solid-media assays (mean ± s.d., *n* = 3 independent replicates). The black dashed line indicates the neutral monoculture baseline (1.0); the red dashed line denotes a 0.5 survival threshold. Asterisks denote pairwise interaction. Significance versus 1.0 was assessed via two-sided one-sample *t*-tests (Supplementary Table 4). Raw CFU ml^-1^ counts are available via Zenodo (DOI: 10.5281/zenodo.23126602).

**Extended Data Fig. 3.**
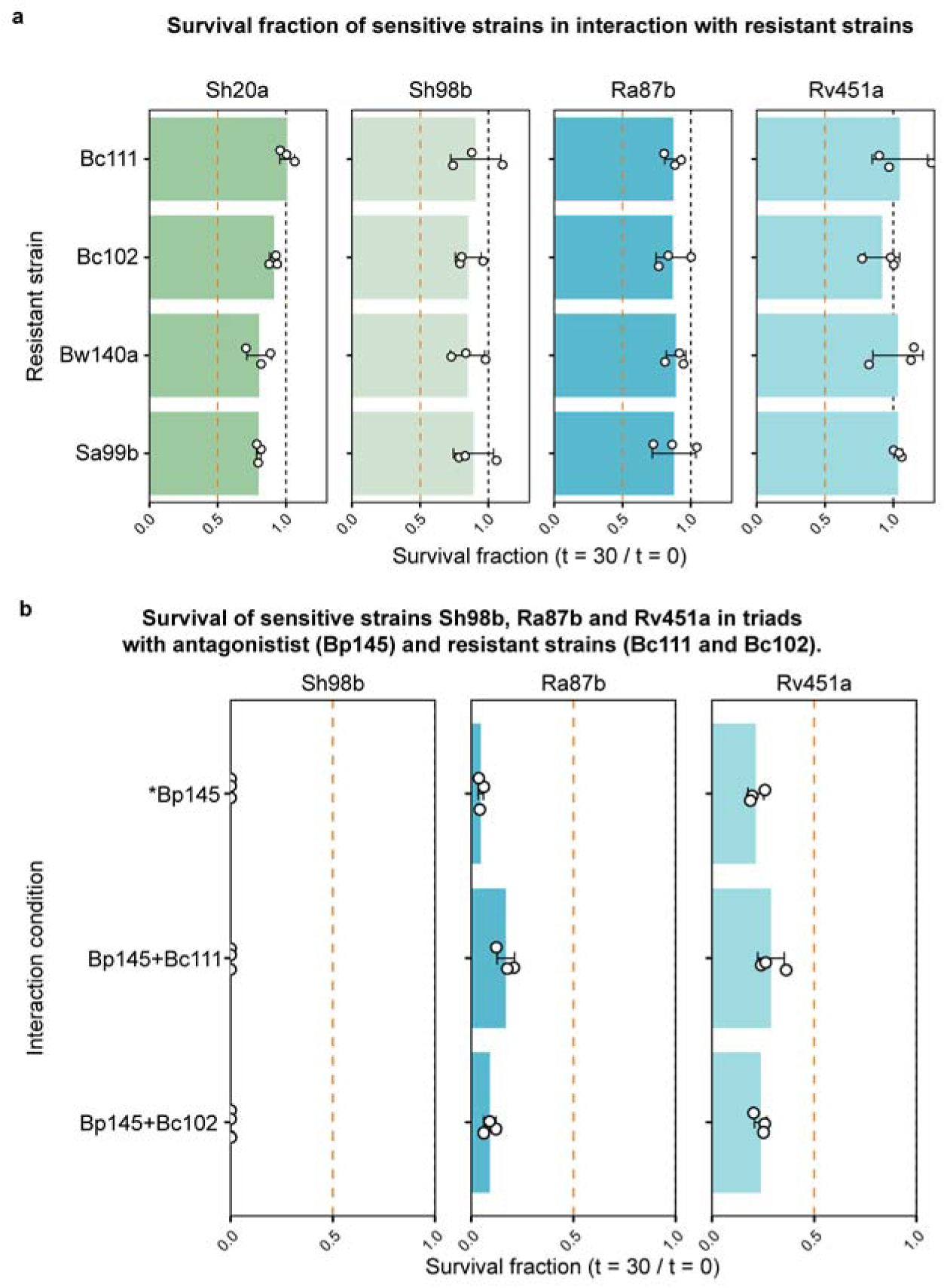
| Baseline neutrality of resistant strains and strain-specific failure of higher-order rescue. **a**, Pairwise survival fractions of sensitive targets co-cultured with candidate resistant strains, confirming the absence of independent antagonistic effects and establishing a neutral pairwise baseline (Supplementary Table 6). **b**, Survival fractions of sensitive strains (Sh98b, Ra87b, and Rv451a) co-cultured with the antagonist *B. pumilus* 145 (Bp145) alone or in triads with resistant strains (Bc111 or Bc102). For all panels, survival fractions were calculated as the ratio of colony-forming units (CFU mL⁻¹) after 30 min of co-culture relative to the initial monoculture (*t*_30_/*t*_0_). Bars represent mean ± s.d. ( independent biological replicates). Dashed lines indicate reference survival thresholds of 1.0 (black; neutral monoculture baseline) and 0.5 (red; strong inhibition). Asterisk denotes pairwise interaction. Significance was assessed via two-sided one- or two-sample *t*-tests (Supplementary Table 7). Raw CFU ml^-1^ counts are available via Zenodo (DOI: 10.5281/zenodo.23126602).

## References

Aguilar-Salinas, B., & Olmedo-Álvarez, G. (2023). A three-species synthetic community model whose rapid response to antagonism allows the study of higher-order dynamics and emergent properties in minutes. Frontiers in Microbiology, 14. 10.3389/fmicb.2023.1057883

Ashkenazy, H., & Weigel, D. (2025). PanGene-O-Meter: Intra-Species Diversity Based on Gene-Content. 10.1101/2025.09.01.673544

Bairey, E., Kelsic, E. D., & Kishony, R. (2016). High-order species interactions shape ecosystem diversity. Nature Communications, 7. 10.1038/ncomms12285

Bankevich, A., Nurk, S., Antipov, D., Gurevich, A. A., Dvorkin, M., Kulikov, A. S., Lesin, V. M., Nikolenko, S. I., Pham, S., Prjibelski, A. D., Pyshkin, A. V., Sirotkin, A. V., Vyahhi, N., Tesler, G., Alekseyev, M. A., & Pevzner, P. A. (2012). SPAdes: A new genome assembly algorithm and its applications to single-cell sequencing. Journal of Computational Biology, 19(5), 455–477. 10.1089/cmb.2012.0021

Blin, K., Shaw, S., Augustijn, H. E., Reitz, Z. L., Biermann, F., Alanjary, M., Fetter, A., Terlouw, B. R., Metcalf, W. W., Helfrich, E. J. N., Van Wezel, G. P., Medema, M. H., & Weber, T. (2023). AntiSMASH 7.0: New and improved predictions for detection, regulation, chemical structures and visualisation. Nucleic Acids Research, 51(W1), W46–W50. 10.1093/nar/gkad344

Casadesús, J., & Low, D. (2006). Epigenetic Gene Regulation in the Bacterial World. Microbiology and Molecular Biology Reviews, 70(3), 830–856. 10.1128/mmbr.00016-06

Chang, C. Y., Bajić, D., Vila, J. C. C., Estrela, S., & Sanchez, A. (2023). Emergent coexistence in multispecies microbial communities. Science, 381(6655), 343–348. 10.1126/science.adg0727

Coyte, K. Z., Schluter, J., & Foster, K. R. (2015). The ecology of the microbiome: Networks, competition, and stability. Science, 350(6261), 663–666. 10.1126/science.aad2602

Eren, A. M., Kiefl, E., Shaiber, A., Veseli, I., Miller, S. E., Schechter, M. S., Fink, I., Pan, J. N., Yousef, M., Fogarty, E. C., Trigodet, F., Watson, A. R., Esen, Ö. C., Moore, R. M., Clayssen, Q., Lee, M. D., Kivenson, V., Graham, E. D., Merrill, B. D., … Willis, A. D. (2020). Community-led, integrated, reproducible multi-omics with anvi’o. In Nature Microbiology (Vol. 6, Number 1, pp. 3–6). Nature Research. 10.1038/s41564-020-00834-3

Faust, K., & Raes, J. (2012). Microbial interactions: From networks to models. In Nature Reviews Microbiology (Vol. 10, Number 8, pp. 538–550). 10.1038/nrmicro2832

Friedman, J., Higgins, L. M., & Gore, J. (2017). Community structure follows simple assembly rules in microbial microcosms. Nature Ecology and Evolution, 1(5). 10.1038/s41559-017-0109

Gallardo-Navarro, O., Arbel-Goren, R., August, E., Olmedo-Alvarez, G., & Stavans, J. (2025). Dynamically induced spatial segregation in multispecies bacterial bioconvection. Nature Communications, 16(1). 10.1038/s41467-025-56244-8

García-García, N., Tamames, J., Linz, A. M., Pedrós-Alió, C., & Puente-Sánchez, F. (2019). Microdiversity ensures the maintenance of functional microbial communities under changing environmental conditions. ISME Journal, 13(12), 2969–2983. 10.1038/s41396-019-0487-8

Ghoul, M., & Mitri, S. (2016). The Ecology and Evolution of Microbial Competition. In Trends in Microbiology (Vol. 24, Number 10, pp. 833–845). Elsevier Ltd. 10.1016/j.tim.2016.06.011

Gómez-Lunar, Z., Hernández-González, I., Rodríguez-Torres, M. D., Souza, V., & Olmedo-álvarez, G. (2016). Microevolution analysis of Bacillus coahuilensis unveils differences in phosphorus acquisition strategies and their regulation. Frontiers in Microbiology, 7(FEB). 10.3389/fmicb.2016.00058

González-Sánchez, A., Cordero-Martínez, E., Rodríguez-Torres, M. D., Islas-Robles, Á., Decena-Segarra, L. P., García-Miranda, N., Navarro-Miranda, M., Zapién-Campos, R., García-Oliva, F., Eguiarte, L. E., Souza, V., & Olmedo-Álvarez, G. (2026). Dormancy as a Signature of Microbial Community Disassembly during Hydrological Collapse in a Desert Oasis. Microbial Ecology. 10.1007/s00248-026-02744-z

González-Torres, P., Pryszcz, L. P., Santos, F., Martínez-García, M., Gabaldón, T., & Antón, J. (2015). Interactions between closely related bacterial strains are revealed by deep transcriptome sequencing. Applied and Environmental Microbiology, 81(24), 8445–8456. 10.1128/AEM.02690-15

Goyal, A., Bittleston, L. S., Leventhal, G. E., Lu, L., & Cordero, O. X. (2022). Interactions between strains govern the eco-evolutionary dynamics of microbial communities. ELife, 11. 10.7554/ELIFE.74987

Grilli, J., Barabás, G., Michalska-Smith, M. J., & Allesina, S. (2017). Higher-order interactions stabilize dynamics in competitive network models. Nature, 548(7666), 210–213. 10.1038/nature23273

Hibbing, M. E., Fuqua, C., Parsek, M. R., & Peterson, S. B. (2010). Bacterial competition: Surviving and thriving in the microbial jungle. In Nature Reviews Microbiology (Vol. 8, Number 1, pp. 15–25). 10.1038/nrmicro2259

Hurtado-Bautista, E., Pérez Sánchez, L. F., Islas-Robles, A., Santoyo, G., & Olmedo-Alvarez, G. (2021). Phenotypic plasticity and evolution of thermal tolerance in bacteria from temperate and hot spring environments. PeerJ, 9. 10.7717/peerj.11734

Jain, C., Rodriguez-R, L. M., Phillippy, A. M., Konstantinidis, K. T., & Aluru, S. (2018). High throughput ANI analysis of 90K prokaryotic genomes reveals clear species boundaries. Nature Communications, 9(1). 10.1038/s41467-018-07641-9

Jia, Y., & Whalen, J. K. (2020). A new perspective on functional redundancy and phylogenetic niche conservatism in soil microbial communities. Pedosphere, 30(1), 18–24. 10.1016/S1002-0160(19)60826-X

Kamrad, S., Aulakh, S. K., Mozzachiodi, S., Blasche, S., Scheidweiler, D., Basile, A., Guan, R., Bradley, R., van den Berg, N. I., Mülleder, M., Ralser, M., & Patil, K. R. (2026). Interspecies interactions drive bacterial proteome reorganization and emergent metabolism. Nature Ecology and Evolution, 10(6), 1127–1144. 10.1038/s41559-026-03030-4

Kelsic, E. D., Zhao, J., Vetsigian, K., & Kishony, R. (2015). Counteraction of antibiotic production and degradation stabilizes microbial communities. Nature, 521(7553), 516–519. 10.1038/nature14485

Kleinhesselink, A. R., Kraft, N. J. B., Pacala, S. W., & Levine, J. M. (2022). Detecting and interpreting higher-order interactions in ecological communities. Ecology Letters, 25(7), 1604–1617. 10.1111/ele.14022

Konstantinidis, K. T., & Tiedje, J. M. (2005). Genomic insights that advance the species definition for prokaryotes. Proceedings of the National Academy of Sciences of the United States of America, 102 (7) 2567-2572. 10.1073/pnas.0409727102

Larkin, A. A., & Martiny, A. C. (2017). Microdiversity shapes the traits, niche space, and biogeography of microbial taxa. In Environmental Microbiology Reports (Vol. 9, Number 2, pp. 55–70). Wiley-Blackwell. 10.1111/1758-2229.12523

Leventhal, G. E., Boix, C., Kuechler, U., Enke, T. N., Sliwerska, E., Holliger, C., & Cordero, O. X. (2018). Strain-level diversity drives alternative community types in millimetre-scale granular biofilms. Nature Microbiology, 3(11), 1295–1303. 10.1038/s41564-018-0242-3

Levine, J. M., Bascompte, J., Adler, P. B., & Allesina, S. (2017). Beyond pairwise mechanisms of species coexistence in complex communities. In Nature (Vol. 546, Number 7656, pp. 56–64). Nature Publishing Group. 10.1038/nature22898

Losos, J. B. (2008). Phylogenetic niche conservatism, phylogenetic signal and the relationship between phylogenetic relatedness and ecological similarity among species. In Ecology Letters (Vol. 11, Number 10, pp. 995–1003). 10.1111/j.1461-0248.2008.01229.x

Louca, S., Polz, M. F., Mazel, F., Albright, M. B. N., Huber, J. A., O’Connor, M. I., Ackermann, M., Hahn, A. S., Srivastava, D. S., Crowe, S. A., Doebeli, M., & Parfrey, L. W. (2018). Function and functional redundancy in microbial systems. In Nature Ecology and Evolution (Vol. 2, Number 6, pp. 936–943). Nature Publishing Group. 10.1038/s41559-018-0519-1

Malfertheiner, L., Tackmann, J., Matias Rodrigues, J. F., & von Mering, C. (2026). Community conservatism is widespread across microbial phyla and environments. Nature Ecology & Evolution. 10.1038/s41559-025-02957-4

Martiny, J. B. H., Jones, S. E., Lennon, J. T., & Martiny, A. C. (2015). Microbiomes in light of traits: A phylogenetic perspective. In Science (Vol. 350, Number 6261). American Association for the Advancement of Science. 10.1126/science.aac9323

Moraïs, S., Mazor, M., Amit, I., Gerth, P., Trautwein-Schult, A., Maaß, S., Grinshpan, I., Shelly, Y., Levin, L., Becher, D., & Mizrahi, I. (2026). Community context reshapes microbial proteomes and reduces functional overlap. Nature Microbiology, 11(5), 1336–1347. 10.1038/s41564-026-02310-w

Moreno-Letelier, A., Olmedo-Alvarez, G., Eguiarte, L. E., & Souza, V. (2012). Divergence and phylogeny of Firmicutes from the Cuatro Ciénegas Basin, Mexico: A window to an ancient ocean. Astrobiology, 12(7), 674–684. 10.1089/ast.2011.0685

Morin, M. A., Morrison, A. J., Harms, M. J., & Dutton, R. J. (2022). Higher-order interactions shape microbial interactions as microbial community complexity increases. Scientific Reports, 12(1). 10.1038/s41598-022-25303-1

Navarro-Muñoz, J. C., Selem-Mojica, N., Mullowney, M. W., Kautsar, S. A., Tryon, J. H., Parkinson, E. I., De Los Santos, E. L. C., Yeong, M., Cruz-Morales, P., Abubucker, S., Roeters, A., Lokhorst, W., Fernandez-Guerra, A., Cappelini, L. T. D., Goering, A. W., Thomson, R. J., Metcalf, W. W., Kelleher, N. L., Barona-Gomez, F., & Medema, M. H. (2020). A computational framework to explore large-scale biosynthetic diversity. Nature Chemical Biology, 16(1), 60–68. 10.1038/s41589-019-0400-9

Niccum, B. A., Kastman, E. K., Kfoury, N., Robbat, A., & Wolfe, B. E. (2020). Strain-Level Diversity Impacts Cheese Rind Microbiome Assembly and Function. MSystems, 5(3). 10.1128/msystems.00149-20

Oña, L., Shreekar, S. K., & Kost, C. (2025). Disentangling microbial interaction networks. In Trends in Microbiology (Vol. 33, Number 6, pp. 619–634). Elsevier Ltd. 10.1016/j.tim.2025.01.013

Patnode, M. L., Guruge, J. L., Castillo, J. J., Couture, G. A., Lombard, V., Terrapon, N., Henrissat, B., Lebrilla, C. B., & Gordon, J. I. (2021). Strain-level functional variation in the human gut microbiota based on bacterial binding to artificial food particles. Cell Host and Microbe, 29(4), 664–673.e5. 10.1016/j.chom.2021.01.007

Pérez-Gutiérrez, R. A., López-Ramírez, V., Islas, Á., Alcaraz, L. D., Hernández-González, I., Olivera, B. C. L., Santillán, M., Eguiarte, L. E., Souza, V., Travisano, M., & Olmedo-Alvarez, G. (2013). Antagonism influences assembly of a Bacillus guild in a local community and is depicted as a food-chain network. ISME Journal, 7(3), 487–497. 10.1038/ismej.2012.119

Rodríguez-Torres, M. D., Islas-Robles, Á., Gómez-Lunar, Z., Delaye, L., Hernández-González, I., Souza, V., Travisano, M., & Olmedo-álvarez, G. (2017). Phenotypic microdiversity and phylogenetic signal analysis of traits related to social interaction in Bacillus spp. from sediment communities. Frontiers in Microbiology, 8(JAN). 10.3389/fmicb.2017.00029

Russel, J., Røder, H. L., Madsen, J. S., Burmølle, M., & Sørensen, S. J. (2017). Antagonism correlates with metabolic similarity in diverse bacteria. Proceedings of the National Academy of Sciences of the United States of America, 114(40), 10684–10688. 10.1073/pnas.1706016114

Sanchez-Gorostiaga, A., Bajić, D., Osborne, M. L., Poyatos, J. F., & Sanchez, A. (2019). High-order interactions distort the functional landscape of microbial consortia. PLoS Biology, 17(12). 10.1371/journal.pbio.3000550

Seemann, T. (2014). Prokka: Rapid prokaryotic genome annotation. Bioinformatics, 30(14), 2068–2069. 10.1093/bioinformatics/btu153

Souza, V., Olmedo-Álvarez, G., & Eguiarte, L. E. (2018). Cuatro Ciénegas Ecology, Natural History and Microbiology. 10.1007/978-3-319-93423-5

Swain, A., Fussell, L., & Fagan, W. F. (2021). Higher-order effects, continuous species interactions, and trait evolution shape microbial spatial dynamics. Proceedings of the National Academy of Sciences, 119(1), e2020956119. 10.1073/pnas.2020956119

Valiente-Banuet, A., Aizen, M. A., Alcántara, J. M., Arroyo, J., Cocucci, A., Galetti, M., García, M. B., García, D., Gómez, J. M., Jordano, P., Medel, R., Navarro, L., Obeso, J. R., Oviedo, R., Ramírez, N., Rey, P. J., Traveset, A., Verdú, M., & Zamora, R. (2015). Beyond species loss: The extinction of ecological interactions in a changing world. Functional Ecology, 29(3), 299–307. 10.1111/1365-2435.12356

van den Berg, N. I., Machado, D., Santos, S., Rocha, I., Chacón, J., Harcombe, W., Mitri, S., & Patil, K. R. (2022). Ecological modelling approaches for predicting emergent properties in microbial communities. In Nature Ecology and Evolution (Vol. 6, Number 7, pp. 855–865). Nature Research. 10.1038/s41559-022-01746-7

Widder, S., Allen, R. J., Pfeiffer, T., Curtis, T. P., Wiuf, C., Sloan, W. T., Cordero, O. X., Brown, S. P., Momeni, B., Shou, W., Kettle, H., Flint, H. J., Haas, A. F., Laroche, B., Kreft, J. U., Rainey, P. B., Freilich, S., Schuster, S., Milferstedt, K., … Wilmes, P. (2016). Challenges in microbial ecology: Building predictive understanding of community function and dynamics. In ISME Journal (Vol. 10, Number 11, pp. 2557–2568). Springer Nature. 10.1038/ismej.2016.45

Wiens, J. J., Ackerly, D. D., Allen, A. P., Anacker, B. L., Buckley, L. B., Cornell, H. V., Damschen, E. I., Jonathan Davies, T., Grytnes, J. A., Harrison, S. P., Hawkins, B. A., Holt, R. D., McCain, C. M., & Stephens, P. R. (2010). Niche conservatism as an emerging principle in ecology and conservation biology. In Ecology Letters (Vol. 13, Number 10, pp. 1310–1324). 10.1111/j.1461-0248.2010.01515.x

Xia, L., Miao, Y., Cao, A., Liu, Y., Liu, Z., Sun, X., Xue, Y., Xu, Z., Xun, W., Shen, Q., Zhang, N., & Zhang, R. (2022). Biosynthetic gene cluster profiling predicts the positive association between antagonism and phylogeny in Bacillus. Nature Communications, 13(1). 10.1038/s41467-022-28668-z

Zapién-Campos, R., Olmedo-álvarez, G., & Santillán, M. (2015). Antagonistic interactions are sufficient to explain self-assemblage of bacterial communities in a homogeneous environment: A computational modeling approach. Frontiers in Microbiology, 6(MAY). 10.3389/fmicb.2015.00489

