## Supplementary Tables 1 to 9 for "Strain-specific higher-order interactions break phylogenetic conservatism in microbial communities"

**Supplementary Table 1 | Metadata of the bacterial strains used in this study.** The dataset comprises 79 environmental isolates from the Churince system and one external reference strain (*Bacillus subtilis* PY79).

| Family | Specie | Strain | Goup | Bioproject | Biosample | Genome Assembly | Selected for interactions | Isolate name |
|---|---|---|---|---|---|---|---|---|
| Bacillaceae | *Bacillus altitudinis* | 22 | Subtilis Clade | PRJNA1232178 | SAMN47227537 | ASM5010801v1 | No | CH22_1T |
| Bacillaceae | *Bacillus altitudinis* | 23 | Subtilis Clade | PRJNA1232178 | SAMN47227538 | ASM5010799v1 | No | CH23_1T |
| Bacillaceae | *Bacillus altitudinis* | 24 | Subtilis Clade | PRJNA1232178 | SAMN47227539 | ASM5010797v1 | No | CH24_1T |
| Bacillaceae | *Bacillus altitudinis* | 38c | Subtilis Clade | PRJNA1232178 | SAMN47227546 | ASM5010783v1 | No | CH38c_1T |
| Bacillaceae | *Bacillus altitudinis* | 156 | Subtilis Clade | PRJNA561254 | SAMN12612709 | ASM818047v1 | Yes | CH156_5T |
| Bacillaceae | *Bacillus atrophaeus* | 90 | Subtilis Clade | PRJNA484220 | SAMN09761861 | ASM868234v1 | Yes | CH90_3T |
| Bacillaceae | *Bacillus cereus* | 164b | Cereus Clade | PRJNA1232178 | SAMN47227573 | ASM5010727v1 | No | CH164b_5T |
| Bacillaceae | *Bacillus cereus* | 102 | Cereus Clade | PRJNA484220 | SAMN09761865 | ASM868228v1 | Yes | CH102_3T |
| Bacillaceae | *Bacillus inaquosorum* | 109a | Subtilis Clade | PRJNA561254 | SAMN12612711 | ASM818034v1 | No | CH109a_4T |
| Bacillaceae | *Bacillus infantis* | 142 | Bacillus infantis | PRJNA1232178 | SAMN47227558 | ASM5010759v1 | No | CH142_4T |
| Bacillaceae | *Bacillus infantis* | 162 | Bacillus infantis | PRJNA1232178 | SAMN47227571 | ASM5010733v1 | No | CH162_5T |
| Bacillaceae | *Bacillus infantis* | 153a | Bacillus infantis | PRJNA1232178 | SAMN47227562 | ASM5010751v1 | No | CH153a_5T |
| Bacillaceae | *Bacillus infantis* | 155a | Bacillus infantis | PRJNA1232178 | SAMN47227564 | ASM5010747v1 | No | CH155a_5T |

| | | | | | | | | |
|---|---|---|---|---|---|---|---|---|
| Bacillaceae | *Bacillus infantis* | 161d | Bacillus infantis | PRJNA1232178 | SAMN47227570 | ASM5010735v1 | No | CH161d_5T |
| Bacillaceae | *Bacillus infantis* | 163b | Bacillus infantis | PRJNA1232178 | SAMN47227572 | ASM5010731v1 | No | CH163b_5T |
| Bacillaceae | *Bacillus infantis* | 37 | Bacillus infantis | PRJNA561254 | SAMN12612703 | ASM818069v1 | No | CH37_1T |
| Bacillaceae | *Bacillus infantis* | 446 | Bacillus infantis | PRJNA561254 | SAMN12612704 | ASM818044v1 | No | CH446_14T |
| Bacillaceae | *Bacillus mobilis* | 157b | Cereus Clade | PRJNA1232178 | SAMN47227565 | ASM5010745v1 | No | CH157b_5T |
| Bacillaceae | *Bacillus mobilis* | 160c | Cereus Clade | PRJNA1232178 | SAMN47227568 | ASM5010739v1 | No | CH160c_5T |
| Bacillaceae | *Bacillus pumilus* | 144b | Subtilis Clade | PRJNA1232178 | SAMN47227559 | ASM5010757v1 | Yes | CH144b_4T |
| Bacillaceae | *Bacillus pumilus* | 145b | Subtilis Clade | PRJNA1232178 | SAMN47227560 | ASM5010755v1 | No | CH145b_4T |
| Bacillaceae | *Bacillus pumilus* | 145 | Subtilis Clade | PRJNA377620 | SAMN06706381 | ASM343197v1 | Yes | CH145_4T |
| Bacillaceae | *Bacillus pumilus* | 150a | Subtilis Clade | PRJNA377620 | SAMN06706382 | ASM357142v1 | Yes | CH150a_4T |
| Bacillaceae | *Bacillus pumilus* | 112a | Subtilis Clade | PRJNA561254 | SAMN12612707 | ASM824526v1 | Yes | CH112a_4T |
| Bacillaceae | *Bacillus pumilus* | 144a | Subtilis Clade | PRJNA561254 | SAMN12612708 | ASM818045v1 | Yes | CH144a_4T |
| Bacillaceae | *Bacillus pumilus* | 95a | Subtilis Clade | PRJNA561254 | SAMN12612706 | ASM818052v1 | Yes | CH95a_3T |
| Bacillaceae | *Bacillus subtilis* | PY79* | Subtilis Clade | PRJNA484220 | SAMN09761863 | ASM868231v1 | Yes | PY79 |
| Bacillaceae | *Bacillus swezeyi* | 21 | Subtilis Clade | PRJNA484220 | SAMN09761860 | ASM868238v1 | Yes | CH21_1T |
| Bacillaceae | *Bacillus swezeyi* | 43 | Subtilis Clade | PRJNA561254 | SAMN12612710 | ASM818046v1 | No | CH43_1T |
| Bacillaceae | *Bacillus thuringiensis* | 33 | Cereus Clade | PRJNA1232178 | SAMN47227544 | ASM5010787v1 | No | CH33_1T |

| | | | | | | | | |
|---|---|---|---|---|---|---|---|---|
| Bacillaceae | *Bacillus thuringiensis* | 93 | Cereus Clade | PRJNA1232178 | SAMN47227554 | ASM5010767v1 | No | CH93_3T |
| Bacillaceae | *Bacillus thuringiensis* | 159b | Cereus Clade | PRJNA1232178 | SAMN47227567 | ASM5010741v1 | No | CH159b_5T |
| Bacillaceae | *Bacillus thuringiensis* | 26b | Cereus Clade | PRJNA1232178 | SAMN47227541 | ASM5010793v1 | No | CH26b_1T |
| Bacillaceae | *Bacillus thuringiensis* | 449b | Cereus Clade | PRJNA1232178 | SAMN47227579 | ASM5010717v1 | No | CH449b_14T |
| Bacillaceae | *Bacillus thuringiensis* | 452a | Cereus Clade | PRJNA1232178 | SAMN47227583 | ASM5010709v1 | No | CH452a_14T |
| Bacillaceae | *Bacillus thuringiensis* | 452b | Cereus Clade | PRJNA1232178 | SAMN47227582 | ASM5010711v1 | No | CH452b_14T |
| Bacillaceae | *Bacillus thuringiensis* | 111 | Cereus Clade | PRJNA561254 | SAMN12879131 | ASM892381v1 | Yes | CH111_4T |
| Bacillaceae | *Bacillus wiedmannii* | 140a | Cereus Clade | PRJNA561254 | SAMN12879137 | ASM892372v1 | Yes | CH140a_4T |
| Micrococcaceae | *Citricoccus zhacaiensis* | 26a | Order Actinomycetales | PRJNA1232178 | SAMN47227542 | ASM5010791v1 | No | CH26a_1T |
| Mycobacteriaceae | *Corynebacterium halotolerans* | 29 | Order Actinomycetales | PRJNA1232178 | SAMN47227543 | ASM5010789v1 | No | CH29_1T |
| Bacillaceae | *Cytobacillus gottheilii* | 449a1 | Cytobacillus gottheilii | PRJNA1232178 | SAMN47227578 | ASM5010719v1 | No | CH449a1_14T |
| Bacillaceae | *Fictibacillus* | 41b | Fictibacillus | PRJNA1232178 | SAMN47227548 | ASM5010779v1 | No | CH41b_1T |

| | | | | | | | | |
|---|---|---|---|---|---|---|---|---|
| | *nanhaiensis* | | nanhaiensis | | | | | |
| Bacillaceae | *Jeotgalibacillus proteolyticus* | 81a | Jeotgalibacillus spp | PRJNA1232178 | SAMN47227551 | ASM5010773v1 | No | CH81a_3T |
| Micrococcaceae | *Kocuria palustris* | 39a | Order Actinomycetales | PRJNA1232178 | SAMN47227547 | ASM5010781v1 | No | CH39a_1T |
| Bacillaceae | *Metabacillus flavus* | 84 | Metabacillus spp | PRJNA1232178 | SAMN47227552 | ASM5010771v1 | No | CH84_3T |
| Bacillaceae | *Metabacillus flavus* | 113a | Metabacillus spp | PRJNA1232178 | SAMN47227556 | ASM5010763v1 | No | CH113a_4T |
| Bacillaceae | *Metabacillus indicus* | 450 | Metabacillus spp | PRJNA1232178 | SAMN47227580 | ASM5010715v1 | No | CH450_14T |
| Micrococcaceae | *Micrococcus luteus* | 149a | Order Actinomycetales | PRJNA1232178 | SAMN47227561 | ASM5010753v1 | No | CH149a_5T |
| Bacillaceae | *Priestia endophytica* | 148 | Megaterium Clade | PRJNA561254 | SAMN12879130 | ASM892380v1 | No | CH148_4T |
| Bacillaceae | *Priestia megaterium* | 447 | Megaterium Clade | PRJNA561254 | SAMN12612716 | ASM818033v1 | No | CH447_14T |
| Bacillaceae | *Rossellomorea aquimaris* | 160a | Aquimaris Clade | PRJNA1232178 | SAMN47227569 | ASM5010737v1 | No | CH160a_5T |
| Bacillaceae | *Rossellomorea aquimaris* | 19a | Aquimaris Clade | PRJNA1232178 | SAMN47227534 | ASM5010807v1 | No | CH19a_1T |
| Bacillaceae | *Rossellomorea aquimaris* | 19b | Aquimaris Clade | PRJNA1232178 | SAMN47227535 | ASM5010805v1 | No | CH19b_1T |
| Bacillaceae | *Rossellomorea aquimaris* | 20b | Aquimaris Clade | PRJNA1232178 | SAMN47227536 | ASM5010803v1 | No | CH20b_1T |

| | | | | | | | | |
|---|---|---|---|---|---|---|---|---|
| Bacillaceae | *Rossellomorea aquimaris* | 30 | Aquimaris Clade | PRJNA560961 | SAMN12604083 | ASM836476v1 | No | CH30_1T |
| Bacillaceae | *Rossellomorea aquimaris* | 159a | Aquimaris Clade | PRJNA561254 | SAMN12612697 | ASM818085v1 | No | CH159b_5T |
| Bacillaceae | *Rossellomorea aquimaris* | 451a | Aquimaris Clade | PRJNA561254 | SAMN12612699 | ASM818077v1 | Yes | CH451a_14T |
| Bacillaceae | *Rossellomorea aquimaris* | 87b | Aquimaris Clade | PRJNA561254 | SAMN12612696 | ASM818081v1 | Yes | CH87b_3T |
| Bacillaceae | *Rossellomorea marisflavi* | 112b | Aquimaris Clade | PRJNA1232178 | SAMN47227555 | ASM5010765v1 | No | CH112b_4T |
| Bacillaceae | *Rossellomorea vietnamensis* | 25 | Aquimaris Clade | PRJNA1232178 | SAMN47227540 | ASM5010795v1 | No | CH25_1T |
| Bacillaceae | *Rossellomorea vietnamensis* | 36 | Aquimaris Clade | PRJNA1232178 | SAMN47227545 | ASM5010785v1 | No | CH36_1T |
| Bacillaceae | *Rossellomorea vietnamensis* | 451b | Aquimaris Clade | PRJNA1232178 | SAMN47227581 | ASM5010713v1 | No | CH451b_14T |
| Bacillaceae | *Rossellomorea vietnamensis* | 138 | Aquimaris Clade | PRJNA377620 | SAMN06706383 | ASM5037945v1 | No | CH138a_1T |
| Bacillaceae | *Rossellomorea vietnamensis* | 34 | Aquimaris Clade | PRJNA561254 | SAMN12612713 | ASM818041v1 | No | CH34_1T |
| Bacillaceae | *Rossellomorea* | 40 | Aquimaris Clade | PRJNA561254 | SAMN12612714 | ASM818061v1 | No | CH40_1T |

| | | | | | | | | |
|---|---|---|---|---|---|---|---|---|
| | *vietnamensis* | | | | | | | |
| Staphylococcaceae | *Staphylococcus arlettae* | 99b | Staphylococcus spp | PRJNA575184 | SAMN12879218 | ASM892333v1 | Yes | CH99b_3T |
| Staphylococcaceae | *Staphylococcus shinii* | 154a | Staphylococcus spp | PRJNA1232178 | SAMN47227563 | ASM5010749v1 | No | CH154a_5T |
| Bacillaceae | *Sutcliffiella halmapala* | 445 | Cohnii Clade | PRJNA1232178 | SAMN47227574 | ASM5010729v1 | No | CH445_14T |
| Bacillaceae | *Sutcliffiella horikoshii* | 44 | Cohnii Clade | PRJNA1232178 | SAMN47227549 | ASM5010777v1 | No | CH44_1T |
| Bacillaceae | *Sutcliffiella horikoshii* | 45 | Cohnii Clade | PRJNA1232178 | SAMN47227550 | ASM5010775v1 | No | CH45_1T |
| Bacillaceae | *Sutcliffiella horikoshii* | 135a | Cohnii Clade | PRJNA1232178 | SAMN47227557 | ASM5010761v1 | No | CH135a_4T |
| Bacillaceae | *Sutcliffiella horikoshii* | 158b | Cohnii Clade | PRJNA1232178 | SAMN47227566 | ASM5010743v1 | No | CH158b_5T |
| Bacillaceae | *Sutcliffiella horikoshii* | 448a | Cohnii Clade | PRJNA1232178 | SAMN47227575 | ASM5010725v1 | No | CH448a_14T |
| Bacillaceae | *Sutcliffiella horikoshii* | 448b | Cohnii Clade | PRJNA1232178 | SAMN47227576 | ASM5010723v1 | No | CH448b_14T |
| Bacillaceae | *Sutcliffiella horikoshii* | 449a2 | Cohnii Clade | PRJNA1232178 | SAMN47227577 | ASM5010721v1 | No | CH449a2_14T |
| Bacillaceae | *Sutcliffiella horikoshii* | 91b | Cohnii Clade | PRJNA1232178 | SAMN47227553 | ASM5010769v1 | No | CH91b_3T |

| | | | | | | | | |
|---|---|---|---|---|---|---|---|---|
| Bacillaceae | *Sutcliffiella horikoshii* | 20a | Cohnii Clade | PRJNA377620 | SAMN06463911 | ASM215785v1 | Yes | CH20a_1T |
| Bacillaceae | *Sutcliffiella horikoshii* | 28 | Cohnii Clade | PRJNA561254 | SAMN12612700 | ASM818072v1 | No | CH28_1T |
| Bacillaceae | *Sutcliffiella horikoshii* | 88 | Cohnii Clade | PRJNA561254 | SAMN12612701 | ASM818084v1 | No | CH88_3T |
| Bacillaceae | *Sutcliffiella horikoshii* | 98b | Cohnii Clade | PRJNA561254 | SAMN12612702 | ASM818071v1 | Yes | CH98b_3T |

**Supplementary Table 2 | Interaction index derived from pairwise inhibition in solid media and genomic traits for network topology.** Out-degree and in-degree represent the number of strains inhibited by, and inhibiting, each focal strain, respectively. Derived indices summarize antagonistic and sensitivity profiles for each strain; formulas used for each metric are indicated in the column headers. A empirical threshold of 0.4 to -0.4 of the Interaction Index Rescaled value was taken to define ecological roles.

| **Strain** | **out_degree** | **in_degree** | **Antagonism Index (out_degree / (N-1))** | **Sensitive Index (in_degree / (N-1))** | **Competition Index (AI - SI)** | **Interaction_Index (CI / Max_AI)** | **II_Rescaled (1 to -1)** | **Ecological Role** | **GC (%)** | **CDS** | **tRNAs** | **rRNAs** | **genome_size_bp** |
|---|---|---|---|---|---|---|---|---|---|---|---|---|---|
| *Bacillus pumilus* 95a | 38 | 0 | 0.49 | 0 | 0.49 | 1.0000 | 1.0000 | Antagonistic | 41 | 3742 | 81 | 15 | 3687435 |
| *Bacillus swezeyi* 21 | 38 | 1 | 0.49 | 0.01 | 0.48 | 0.9796 | 0.9759 | Antagonistic | 44 | 4772 | 86 | 13 | 4762017 |

| | | | | | | | | | | | | | |
|---|---|---|---|---|---|---|---|---|---|---|---|---|---|
| *Bacillus atrophaeus* 90 | 36 | 0 | 0.47 | 0 | 0.47 | 0.9592 | 0.9518 | Antagonistic | 43 | 4038 | 82 | 12 | 4136251 |
| *Bacillus altitudinis* 156 | 34 | 0 | 0.44 | 0 | 0.44 | 0.8980 | 0.8795 | Antagonistic | 41 | 3836 | 76 | 18 | 3784785 |
| *Bacillus pumilus* 150a | 34 | 0 | 0.44 | 0 | 0.44 | 0.8980 | 0.8795 | Antagonistic | 41 | 3726 | 82 | 24 | 3747740 |
| *Bacillus swezeyi* 43 | 31 | 0 | 0.4 | 0 | 0.4 | 0.8163 | 0.7831 | Antagonistic | 44 | 4620 | 86 | 15 | 4614405 |
| *Bacillus altitudinis* 22 | 29 | 0 | 0.38 | 0 | 0.38 | 0.7755 | 0.7349 | Antagonistic | 41 | 3953 | 79 | 11 | 3862232 |
| *Bacillus pumilus* 145 | 32 | 3 | 0.42 | 0.04 | 0.38 | 0.7755 | 0.7349 | Antagonistic | 41 | 3983 | 81 | 24 | 3989121 |
| *Bacillus pumilus* 144a | 32 | 4 | 0.42 | 0.05 | 0.37 | 0.7551 | 0.7108 | Antagonistic | 41 | 4129 | 80 | 14 | 4024237 |
| *Bacillus pumilus* 144b | 32 | 4 | 0.42 | 0.05 | 0.37 | 0.7551 | 0.7108 | Antagonistic | 41 | 4094 | 74 | 12 | 3988044 |
| *Bacillus inaquosorum* 109a | 31 | 6 | 0.4 | 0.08 | 0.32 | 0.6531 | 0.5904 | Antagonistic | 44 | 4201 | 86 | 20 | 4260972 |
| *Staphylococcus shinii* 154a | 13 | 0 | 0.17 | 0 | 0.17 | 0.3469 | 0.2289 | Resistant | 32 | 2845 | 60 | 12 | 3027272 |
| *Bacillus infantis* 37 | 10 | 0 | 0.13 | 0 | 0.13 | 0.2653 | 0.1325 | Resistant | 45 | 5241 | 88 | 18 | 5168596 |
| *Metabacillus flavus* 113a | 9 | 0 | 0.12 | 0 | 0.12 | 0.2449 | 0.1084 | Resistant | 45 | 3841 | 83 | 11 | 3857092 |

| | | | | | | | | | | | | | |
|---|---|---|---|---|---|---|---|---|---|---|---|---|---|
| *Priestia endophyticus* 148 | 10 | 1 | 0.13 | 0.01 | 0.12 | 0.2449 | 0.1084 | Resistant | 37 | 5317 | 82 | 12 | 5346832 |
| *Bacillus altitudinis* 23 | 8 | 1 | 0.1 | 0.01 | 0.09 | 0.1837 | 0.0362 | Resistant | 41 | 3955 | 79 | 12 | 3863395 |
| *Rossellomorea marisflavi* 112b | 10 | 3 | 0.13 | 0.04 | 0.09 | 0.1837 | 0.0362 | Resistant | 48 | 4588 | 105 | 13 | 4477587 |
| *Sutcliffiella horikoshii* 448b | 7 | 0 | 0.09 | 0 | 0.09 | 0.1837 | 0.0362 | Resistant | 41 | 4324 | 81 | 15 | 4262389 |
| *Rossellomorea aquimaris* 20b | 8 | 2 | 0.1 | 0.03 | 0.07 | 0.1429 | -0.0120 | Resistant | 41 | 4676 | 109 | 16 | 4735417 |
| *Metabacillus indicus* 450 | 8 | 3 | 0.1 | 0.04 | 0.06 | 0.1224 | -0.0361 | Resistant | 44 | 4148 | 82 | 17 | 4073752 |
| *Rossellomorea aquimaris* 30 | 5 | 0 | 0.06 | 0 | 0.06 | 0.1224 | -0.0361 | Resistant | 41 | 4856 | 109 | 15 | 4912469 |
| *Bacillus thuringiensis* 449b | 3 | 0 | 0.04 | 0 | 0.04 | 0.0816 | -0.0843 | Resistant | 35 | 6058 | 101 | 21 | 6009662 |
| *Fictibacillus nanhaiensis* 41b | 6 | 3 | 0.08 | 0.04 | 0.04 | 0.0816 | -0.0843 | Resistant | 39 | 4278 | 68 | 12 | 4221762 |
| *Bacillus altitudinis* 24 | 2 | 0 | 0.03 | 0 | 0.03 | 0.0612 | -0.1084 | Resistant | 41 | 3956 | 79 | 12 | 3863983 |
| *Rossellomorea* | 2 | 0 | 0.03 | 0 | 0.03 | 0.0612 | -0.1084 | Resistant | 41 | 4673 | 108 | 16 | 4734122 |

| | | | | | | | | | | | | | |
|---|---|---|---|---|---|---|---|---|---|---|---|---|---|
| *aquimaris* 19b | | | | | | | | | | | | | |
| *Bacillus altitudinis* 38c | 1 | 0 | 0.01 | 0 | 0.01 | 0.0204 | -0.1566 | Resistant | 41 | 3836 | 78 | 18 | 3785011 |
| *Bacillus cereus* 164b | 1 | 0 | 0.01 | 0 | 0.01 | 0.0204 | -0.1566 | Resistant | 35 | 6051 | 104 | 18 | 6028049 |
| *Bacillus thuringiensis* 159b | 1 | 0 | 0.01 | 0 | 0.01 | 0.0204 | -0.1566 | Resistant | 35 | 5970 | 102 | 20 | 5910033 |
| *Bacillus thuringiensis* 452b | 1 | 0 | 0.01 | 0 | 0.01 | 0.0204 | -0.1566 | Resistant | 35 | 6061 | 104 | 21 | 6042849 |
| *Bacillus pumilus* 112a | 0 | 0 | 0 | 0 | 0 | 0.0000 | -0.1807 | Resistant | 41 | 3745 | 83 | 14 | 3688733 |
| *Bacillus thuringiensis* 452a | 14 | 14 | 0.18 | 0.18 | 0 | 0.0000 | -0.1807 | Resistant | 35 | 5975 | 108 | 17 | 5956691 |
| *Kocuria palustris* 39a | 0 | 0 | 0 | 0 | 0 | 0.0000 | -0.1807 | Resistant | 70 | 2587 | 51 | 7 | 2948183 |
| *Staphylococcus arlettae* 99b | 0 | 0 | 0 | 0 | 0 | 0.0000 | -0.1807 | Resistant | 33 | 2435 | 60 | 10 | 2580826 |
| *Sutcliffiella horikoshii* 448a | 0 | 0 | 0 | 0 | 0 | 0.0000 | -0.1807 | Resistant | 41 | 4322 | 81 | 15 | 4263539 |
| *Sutcliffiella horikoshii* 88 | 0 | 0 | 0 | 0 | 0 | 0.0000 | -0.1807 | Resistant | 41 | 4149 | 81 | 14 | 4124408 |
| *Bacillus thuringiensis* 93 | 0 | 1 | 0 | 0.01 | -0.01 | -0.0204 | -0.2048 | Resistant | 35 | 6016 | 95 | 17 | 6157038 |

| | | | | | | | | | | | | |
|---|---|---|---|---|---|---|---|---|---|---|---|---|
| *Bacillus thuringiensis* 26b | 0 | 2 | 0 | 0.03 | -0.03 | -0.0612 | -0.2530 | Resistant | 35 | 5967 | 99 | 17 | 5962055 |
| *Corynebacterium halotolerans* 29 | 0 | 2 | 0 | 0.03 | -0.03 | -0.0612 | -0.2530 | Resistant | 66 | 3008 | 56 | 6 | 3338894 |
| *Bacillus mobilis* 160c | 0 | 3 | 0 | 0.04 | -0.04 | -0.0816 | -0.2771 | Resistant | 35 | 5559 | 109 | 20 | 5541725 |
| *Rossellomorea aquimaris* 87b | 8 | 11 | 0.1 | 0.14 | -0.04 | -0.0816 | -0.2771 | Resistant | 41 | 4726 | 100 | 13 | 4753727 |
| *Bacillus thuringiensis* 111 | 0 | 4 | 0 | 0.05 | -0.05 | -0.1020 | -0.3012 | Resistant | 35 | 5917 | 92 | 16 | 5965683 |
| *Bacillus wiedmannii* 140a | 0 | 4 | 0 | 0.05 | -0.05 | -0.1020 | -0.3012 | Resistant | 35 | 5506 | 90 | 11 | 5376841 |
| *Rossellomorea vietnamensis* 25 | 0 | 4 | 0 | 0.05 | -0.05 | -0.1020 | -0.3012 | Resistant | 42 | 4566 | 85 | 16 | 4821045 |
| *Bacillus mobilis* 157b | 0 | 5 | 0 | 0.06 | -0.06 | -0.1224 | -0.3253 | Resistant | 35 | 5533 | 106 | 17 | 5525117 |
| *Sutcliffiella horikoshii* 45 | 0 | 6 | 0 | 0.08 | -0.08 | -0.1633 | -0.3735 | Resistant | 41 | 4266 | 81 | 14 | 4208236 |
| *Cytobacillus gottheilii* 449a1 | 0 | 7 | 0 | 0.09 | -0.09 | -0.1837 | -0.3976 | Resistant | 39 | 4173 | 78 | 22 | 4320919 |
| *Rossellomorea vietnamensis* 36 | 1 | 8 | 0.01 | 0.1 | -0.09 | -0.1837 | -0.3976 | Resistant | 42 | 4463 | 84 | 18 | 4650283 |

| | | | | | | | | | | | | | |
|---|---|---|---|---|---|---|---|---|---|---|---|---|---|
| *Bacillus infantis* 142 | 2 | 10 | 0.03 | 0.13 | -0.1 | -0.2041 | -0.4217 | Sensitive | 46 | 4984 | 89 | 16 | 4908856 |
| *Jeotgalibacillus proteolyticus* 81a | 0 | 8 | 0 | 0.1 | -0.1 | -0.2041 | -0.4217 | Sensitive | 42 | 3705 | 78 | 12 | 3746204 |
| *Rossellomorea vietnamensis* 451b | 0 | 8 | 0 | 0.1 | -0.1 | -0.2041 | -0.4217 | Sensitive | 42 | 4423 | 84 | 14 | 4629518 |
| *Sutcliffiella horikoshii* 91b | 0 | 8 | 0 | 0.1 | -0.1 | -0.2041 | -0.4217 | Sensitive | 41 | 4200 | 77 | 10 | 4193951 |
| *Rossellomorea aquimaris* 160a | 4 | 12 | 0.05 | 0.16 | -0.11 | -0.2245 | -0.4458 | Sensitive | 41 | 4712 | 105 | 20 | 4730458 |
| *Sutcliffiella horikoshii* 44 | 0 | 9 | 0 | 0.12 | -0.12 | -0.2449 | -0.4699 | Sensitive | 41 | 4265 | 81 | 14 | 4206829 |
| *Bacillus infantis* 155a | 0 | 10 | 0 | 0.13 | -0.13 | -0.2653 | -0.4940 | Sensitive | 46 | 5024 | 90 | 21 | 5002861 |
| *Citricoccus zhacaiensis* 26a | 0 | 10 | 0 | 0.13 | -0.13 | -0.2653 | -0.4940 | Sensitive | 71 | 3005 | 52 | 5 | 3475792 |
| *Micrococcus luteus* 149a | 0 | 10 | 0 | 0.13 | -0.13 | -0.2653 | -0.4940 | Sensitive | 73 | 2283 | 57 | 5 | 2525489 |
| *Rossellomorea aquimaris* 19a | 0 | 10 | 0 | 0.13 | -0.13 | -0.2653 | -0.4940 | Sensitive | 41 | 4673 | 108 | 16 | 4733948 |
| *Bacillus thuringiensis* 33 | 0 | 11 | 0 | 0.14 | -0.14 | -0.2857 | -0.5181 | Sensitive | 35 | 5972 | 105 | 17 | 5966767 |

| | | | | | | | | | | | | | |
|---|---|---|---|---|---|---|---|---|---|---|---|---|---|
| *Metabacillus flavus* 84 | 0 | 11 | 0 | 0.14 | -0.14 | -0.2857 | -0.5181 | Sensitive | 45 | 3948 | 84 | 15 | 3932029 |
| *Sutcliffiella horikoshii* 135a | 0 | 11 | 0 | 0.14 | -0.14 | -0.2857 | -0.5181 | Sensitive | 41 | 4203 | 83 | 15 | 4203282 |
| *Sutcliffiella horikoshii* 28 | 0 | 11 | 0 | 0.14 | -0.14 | -0.2857 | -0.5181 | Sensitive | 40 | 4559 | 81 | 12 | 4522185 |
| *Sutcliffiella horikoshii* 98b | 0 | 11 | 0 | 0.14 | -0.14 | -0.2857 | -0.5181 | Sensitive | 41 | 4207 | 82 | 15 | 4209183 |
| *Priestia megaterium* 447 | 0 | 12 | 0 | 0.16 | -0.16 | -0.3265 | -0.5662 | Sensitive | 37 | 4066 | 98 | 16 | 4081202 |
| *Bacillus infantis* 161d | 1 | 14 | 0.01 | 0.18 | -0.17 | -0.3469 | -0.5903 | Sensitive | 46 | 5126 | 90 | 17 | 5108662 |
| *Rossellomorea vietnamensis* 138 | 0 | 13 | 0 | 0.17 | -0.17 | -0.3469 | -0.5903 | Sensitive | 42 | 4591 | 84 | 33 | 4808993 |
| *Sutcliffiella horikoshii* 158b | 0 | 13 | 0 | 0.17 | -0.17 | -0.3469 | -0.5903 | Sensitive | 40 | 4342 | 82 | 11 | 4282248 |
| *Rossellomorea aquimaris* 159a | 3 | 17 | 0.04 | 0.22 | -0.18 | -0.3673 | -0.6144 | Sensitive | 42 | 4718 | 113 | 14 | 4744327 |
| *Sutcliffiella horikoshii* 20a | 0 | 14 | 0 | 0.18 | -0.18 | -0.3673 | -0.6144 | Sensitive | 41 | 4292 | 74 | 24 | 4277585 |
| *Sutcliffiella horikoshii* 449a2 | 0 | 14 | 0 | 0.18 | -0.18 | -0.3673 | -0.6144 | Sensitive | 41 | 4325 | 81 | 15 | 4262321 |

| | | | | | | | | | | | | |
|---|---|---|---|---|---|---|---|---|---|---|---|---|
| *Sutcliffiella halmapalus* 445 | 0 | 15 | 0 | 0.19 | -0.19 | -0.3878 | -0.6385 | Sensitive | 42 | 4519 | 82 | 14 | 4444703 |
| *Bacillus infantis* 162 | 1 | 16 | 0.01 | 0.21 | -0.2 | -0.4082 | -0.6626 | Sensitive | 46 | 5114 | 90 | 17 | 5098514 |
| *Bacillus pumilus* 145b | 1 | 16 | 0.01 | 0.21 | -0.2 | -0.4082 | -0.6626 | Sensitive | 41 | 4097 | 74 | 12 | 3937399 |
| *Bacillus infantis* 446 | 1 | 17 | 0.01 | 0.22 | -0.21 | -0.4286 | -0.6867 | Sensitive | 46 | 5151 | 89 | 25 | 5144535 |
| *Rossellomorea vietnamensis* 40 | 0 | 16 | 0 | 0.21 | -0.21 | -0.4286 | -0.6867 | Sensitive | 42 | 4522 | 82 | 20 | 4704930 |
| *Bacillus infantis* 163b | 0 | 17 | 0 | 0.22 | -0.22 | -0.4490 | -0.7108 | Sensitive | 45 | 5334 | 87 | 21 | 5167900 |
| *Rossellomorea vietnamensis* 451a | 0 | 17 | 0 | 0.22 | -0.22 | -0.4490 | -0.7108 | Sensitive | 42 | 4439 | 84 | 14 | 4656734 |
| *Bacillus infantis* 153a | 0 | 19 | 0 | 0.25 | -0.25 | -0.5102 | -0.7831 | Sensitive | 46 | 5100 | 88 | 19 | 5038092 |
| *Rossellomorea vietnamensis* 34 | 0 | 26 | 0 | 0.34 | -0.34 | -0.6939 | -1.0000 | Sensitive | 42 | 4585 | 82 | 19 | 4817581 |

**Supplementary Table 3 | Reference genomes used in the pangenome analysis.** Reference genomes were selected based on RefSeq annotation quality from NCBI at the time of analysis. For *Sutcliffiella horikoshii*, an alternative high-quality RefSeq strain was used as the external reference, given that *S. horikoshii* 20a belongs to the local environmental collection evaluated in this study.

| RefSeq Genome | NCBI RefSeq | Biosample | Strain |
|---|---|---|---|
| *Bacillus altitudinis* | GCF_001191605.1 | SAMN02905161 | GR-8 |
| *Bacillus atrophaeus* | GCF_039519175.1 | SAMN41078598 | SW |
| *Bacillus cereus* | GCF_006094295.1 | SAMN10591533 | ATCC 14579 |
| *Bacillus inaquosorum* | GCF_023238325.1 | SAMN27761609 | A65.1 |
| *Bacillus infantis* | GCF_979912685.1 | SAMEA11878141 5 | SRL369 |
| *Bacillus mobilis* | GCF_057198225.1 | SAMN47812969 | LaCa_104 |
| *Bacillus pumilus* | GCF_000017885.4 | SAMN00253833 | SAFR-032 |
| *Bacillus subtilis* | GCF_000009045.1 | SAMEA3138188 | 168 |
| *Bacillus swezeyi* | GCF_030908325.1 | SAMN37015269 | EA208 |
| *Bacillus thuringiensis* | GCF_000161615.1 | SAMN00738287 | ATCC 10792 |
| *Bacillus wiedmannii* | GCF_055414345.1 | SAMN43997841 | NPDC092377 |
| *Citricoccus zhacaiensis* | GCF_014645315.1 | SAMD00245148 | CGMCC 1.7064 |
| *Corynebacterium halotolerans* | GCF_000341345.1 | SAMN02603027 | YIM 70093 |
| *Cytobacillus gottheilii* | GCF_018582665.1 | SAMN18214881 | 1839 |
| *Fictibacillus nanhaiensis* | GCF_042686385.1 | SAMN43282316 | KCTC 13712 |
| *Jeotgalibacillus campisalis* | GCF_000829515.1 | SAMN03009544 | 22-7 22-7 |
| *Kocuria palustris* | GCF_016907795.1 | SAMN17619998 | TAGA27 |
| *Metabacillus indicus* | GCF_046603695.1 | SAMN45866069 | JKLC.Or24 |
| *Metabacillus flavus* | GCF_018283675.1 | SAMN18642694 | KIGAM252 |
| *Micrococcus luteus* | GCF_900475555.1 | SAMEA3867442 | NCTC2665 |
| *Priestia endophytica* | GCF_900115845.1 | SAMN02745910 | DSM 13796 |
| *Priestia megaterium* | GCF_006094495.1 | SAMN10738403 | ATCC 14581 |
| *Rossellomorea vietnamensis* | GCF_000723905.1 | SAMEA3139053 | HD-02 |
| *Rossellomorea aquimaris* | GCF_001648555.1 | SAMN04403367 | TF12 |
| *Rossellomorea marisflavi* | GCF_022170785.1 | SAMN20890003 | UTDF19-31A |
| *Staphylococcus arlettae* | GCF_025560845.1 | SAMN26923316 | IVB6235 |
| *Staphylococcus shinii* | GCF_017583065.1 | SAMN18334909 | K22-5M |
| *Sutcliffiella halmapala* | GCF_002019665.1 | SAMN05934582 | DSM 8723 |
| *Sutcliffiella horikoshii* | GCF_044809035.1 | SAMN22138092 | ADH2-198 |

**Supplementary Table 4 | Pairwise interaction outcomes of sensitive strains.** Survival fractions of sensitive targets following co-culture with individual antagonistic or candidate resistant strains. Data are presented as mean ± s.d. with 95% confidence intervals (CI) from three independent biological replicates (n = 3). Statistical significance was assessed against a neutral monoculture baseline (survival fraction = 1.0) using two-sided one-sample t-tests. P values were adjusted for multiple comparisons using the Benjamini–Hochberg procedure, with significance determined by the resulting adjusted P values (q).

| **Antagonist** | **Sensitive** | **N** | **Survivor fraction mean of S** | **SD** | **CI95_low** | **CI95_high** | **p_raw** | **p_adj** | **Significance** | **Mean ± SD (CI95%)** |
|---|---|---|---|---|---|---|---|---|---|---|
| *Bacillus pumilus* 145 | *Rossellomorea aquimaris* 87b | 3 | 0.814 | 0.018 | 0.793 | 0.835 | 0.003 | 0.014 | * | 0.814 ± 0.018 (0.793–0.835) |
| *Bacillus pumilus* 145 | *Rossellomorea vietnamensis* 451a | 3 | 0.056 | 0.005 | 0.050 | 0.063 | 0.000 | 0.000 | *** | 0.056 ± 0.005 (0.050–0.063) |
| *Bacillus pumilus* 145 | *Sutcliffiella horikoshii* 20a | 3 | 0.206 | 0.067 | 0.129 | 0.282 | 0.002 | 0.013 | * | 0.206 ± 0.067 (0.129–0.282) |
| *Bacillus pumilus* 145 | *Sutcliffiella horikoshii* 98b | 3 | 0.741 | 0.023 | 0.715 | 0.767 | 0.003 | 0.014 | * | 0.741 ± 0.023 (0.715–0.767) |
| *Bacillus pumilus* 95a | *Rossellomorea aquimaris* 87b | 3 | 0.374 | 0.158 | 0.194 | 0.553 | 0.021 | 0.058 | | 0.374 ± 0.158 (0.194–0.553) |
| *Bacillus pumilus* 95a | *Rossellomorea vietnamensis* 451a | 3 | 0.470 | 0.132 | 0.321 | 0.619 | 0.020 | 0.058 | | 0.470 ± 0.132 (0.321–0.619) |

| | | | | | | | | | | |
|---|---|---|---|---|---|---|---|---|---|---|
| *Bacillus pumilus* 95a | *Sutcliffiella horikoshii* 20a | 3 | 0.326 | 0.085 | 0.230 | 0.422 | 0.005 | 0.020 | * | 0.326 ± 0.085 (0.230–0.422) |
| *Bacillus pumilus* 95a | *Sutcliffiella horikoshii* 98b | 3 | 0.018 | 0.008 | 0.009 | 0.026 | 0.000 | 0.000 | *** | 0.018 ± 0.008 (0.009–0.026) |
| *Bacillus pumilus* 144b | *Rossellomorea aquimaris* 87b | 3 | 0.526 | 0.078 | 0.438 | 0.614 | 0.009 | 0.029 | * | 0.526 ± 0.078 (0.438–0.614) |
| *Bacillus pumilus* 144b | *Rossellomorea vietnamensis* 451a | 3 | 0.001 | 0.001 | -0.001 | 0.002 | 0.000 | 0.000 | *** | 0.001 ± 0.001 (-0.001–0.002) |
| *Bacillus pumilus* 144b | *Sutcliffiella horikoshii* 20a | 3 | 0.359 | 0.158 | 0.180 | 0.537 | 0.020 | 0.058 | | 0.359 ± 0.158 (0.180–0.537) |
| *Bacillus pumilus* 144b | *Sutcliffiella horikoshii* 98b | 3 | 0.656 | 0.209 | 0.419 | 0.892 | 0.104 | 0.188 | | 0.656 ± 0.209 (0.419–0.892) |
| *Bacillus pumilus* 112a | *Rossellomorea aquimaris* 87b | 3 | 0.598 | 0.156 | 0.421 | 0.775 | 0.047 | 0.105 | | 0.598 ± 0.156 (0.421–0.775) |
| *Bacillus pumilus* 112a | *Rossellomorea vietnamensis* 451a | 3 | 0.001 | 0.001 | -0.001 | 0.002 | 0.000 | 0.000 | *** | 0.001 ± 0.001 (-0.001–0.002) |
| *Bacillus pumilus* 112a | *Sutcliffiella horikoshii* 20a | 3 | 0.409 | 0.074 | 0.325 | 0.493 | 0.005 | 0.020 | * | 0.409 ± 0.074 (0.325–0.493) |

| | | | | | | | | | | |
|---|---|---|---|---|---|---|---|---|---|---|
| *Bacillus pumilus* 112a | *Sutcliffiella horikoshii* 98b | 3 | 0.021 | 0.006 | 0.014 | 0.028 | 0.000 | 0.000 | *** | 0.021 ± 0.006 (0.014–0.028) |
| *Bacillus pumilus* 150a | *Rossellomorea aquimaris* 87b | 3 | 1.026 | 0.103 | 0.910 | 1.143 | 0.701 | 0.770 | | 1.026 ± 0.103 (0.910–1.143) |
| *Bacillus pumilus* 150a | *Rossellomorea vietnamensis* 451a | 3 | 0.634 | 0.103 | 0.517 | 0.750 | 0.025 | 0.067 | | 0.634 ± 0.103 (0.517–0.750) |
| *Bacillus pumilus* 150a | *Sutcliffiella horikoshii* 20a | 3 | 0.575 | 0.185 | 0.366 | 0.784 | 0.058 | 0.124 | | 0.575 ± 0.185 (0.366–0.784) |
| *Bacillus pumilus* 150a | *Sutcliffiella horikoshii* 98b | 3 | 0.279 | 0.022 | 0.254 | 0.303 | 0.000 | 0.002 | ** | 0.279 ± 0.022 (0.254–0.303) |
| *Bacillus pumilus* 156 | *Rossellomorea aquimaris* 87b | 3 | 1.107 | 0.084 | 1.012 | 1.202 | 0.157 | 0.258 | | 1.107 ± 0.084 (1.012–1.202) |
| *Bacillus pumilus* 156 | *Rossellomorea vietnamensis* 451a | 3 | 0.663 | 0.033 | 0.625 | 0.700 | 0.003 | 0.014 | * | 0.663 ± 0.033 (0.625–0.700) |
| *Bacillus pumilus* 156 | *Sutcliffiella horikoshii* 20a | 3 | 1.014 | 0.157 | 0.836 | 1.192 | 0.891 | 0.907 | | 1.014 ± 0.157 (0.836–1.192) |
| *Bacillus pumilus* 156 | *Sutcliffiella horikoshii* 98b | 3 | 0.800 | 0.097 | 0.690 | 0.909 | 0.070 | 0.137 | | 0.800 ± 0.097 (0.690–0.909) |

| | | | | | | | | | | |
|---|---|---|---|---|---|---|---|---|---|---|
| *Bacillus swezeyi* 21 | *Rossellomorea aquimaris* 87b | 3 | 0.879 | 0.185 | 0.669 | 1.089 | 0.375 | 0.467 | | 0.879 ± 0.185 (0.669–1.089) |
| *Bacillus swezeyi* 21 | *Rossellomorea vietnamensis* 451a | 3 | 0.942 | 0.180 | 0.738 | 1.146 | 0.632 | 0.708 | | 0.942 ± 0.180 (0.738–1.146) |
| *Bacillus swezeyi* 21 | *Sutcliffiella horikoshii* 20a | 3 | 0.707 | 0.237 | 0.438 | 0.975 | 0.166 | 0.265 | | 0.707 ± 0.237 (0.438–0.975) |
| *Bacillus swezeyi* 21 | *Sutcliffiella horikoshii* 98b | 3 | 1.085 | 0.096 | 0.977 | 1.194 | 0.263 | 0.360 | | 1.085 ± 0.096 (0.977–1.194) |
| *Bacillus atrophaeus* 90 | *Rossellomorea aquimaris* 87b | 3 | 0.888 | 0.214 | 0.645 | 1.130 | 0.459 | 0.536 | | 0.888 ± 0.214 (0.645–1.130) |
| *Bacillus atrophaeus* 90 | *Rossellomorea vietnamensis* 451a | 3 | 0.858 | 0.070 | 0.779 | 0.938 | 0.073 | 0.137 | | 0.858 ± 0.070 (0.779–0.938) |
| *Bacillus atrophaeus* 90 | *Sutcliffiella horikoshii* 20a | 3 | 0.921 | 0.072 | 0.839 | 1.003 | 0.198 | 0.290 | | 0.921 ± 0.072 (0.839–1.003) |
| *Bacillus atrophaeus* 90 | *Sutcliffiella horikoshii* 98b | 3 | 1.062 | 0.108 | 0.940 | 1.184 | 0.422 | 0.503 | | 1.062 ± 0.108 (0.940–1.184) |
| *Bacillus pumilus* 144a | *Rossellomorea aquimaris* 87b | 3 | 0.653 | 0.102 | 0.539 | 0.768 | 0.027 | 0.070 | | 0.653 ± 0.102 (0.539–0.768) |

| | | | | | | | | | | |
|---|---|---|---|---|---|---|---|---|---|---|
| *Bacillus pumilus* 144a | *Rossellomorea vietnamensis* 451a | 3 | 0.016 | 0.008 | 0.006 | 0.025 | 0.000 | 0.000 | *** | 0.016 ± 0.008 (0.006–0.025) |
| *Bacillus pumilus* 144a | *Sutcliffiella horikoshii* 20a | 3 | 0.709 | 0.103 | 0.593 | 0.825 | 0.039 | 0.095 | | 0.709 ± 0.103 (0.593–0.825) |
| *Bacillus pumilus* 144a | *Sutcliffiella horikoshii* 98b | 3 | 0.514 | 0.069 | 0.436 | 0.591 | 0.007 | 0.023 | * | 0.514 ± 0.069 (0.436–0.591) |
| *Bacillus subtilis* PY79 | *Rossellomorea aquimaris* 87b | 3 | 0.887 | 0.146 | 0.722 | 1.052 | 0.310 | 0.410 | | 0.887 ± 0.146 (0.722–1.052) |
| *Bacillus subtilis* PY79 | *Rossellomorea vietnamensis* 451a | 3 | 0.026 | 0.026 | -0.003 | 0.055 | 0.000 | 0.002 | ** | 0.026 ± 0.026 (-0.003–0.055) |
| *Bacillus subtilis* PY79 | *Sutcliffiella horikoshii* 20a | 3 | 0.830 | 0.174 | 0.633 | 1.027 | 0.234 | 0.327 | | 0.830 ± 0.174 (0.633–1.027) |
| *Bacillus subtilis* PY79 | *Sutcliffiella horikoshii* 98b | 3 | 0.995 | 0.095 | 0.888 | 1.102 | 0.932 | 0.932 | | 0.995 ± 0.095 (0.888–1.102) |

**Supplementary Table 5 | Statistical assessment of strain-specific antagonistic activity.** Survival fractions ($t_{30}/t_0$) were tested against a neutral monoculture baseline (survival fraction = 1.0) using two-sided Wilcoxon signed-rank tests. The table reports sample size (n), median survival, test statistic (W), nominal P values, and Benjamini–Hochberg adjusted P values (q). Interactions with q < 0.05 were considered statistically significant. Individual survival fraction replicates are provided in the Source Data.

| Antagonistic Strain | Survival fraction of Sensitive Strains Median | N | p_raw | Statistic | p_adj | Significant |
|---|---|---|---|---|---|---|
| *Bacillus pumilus* 112a | 0.17712 | 12 | 0.00049 | 0 | 0.00098 | True |
| *Bacillus pumilus* 95a | 0.33562 | 12 | 0.00049 | 0 | 0.00098 | True |
| *Bacillus pumilus* 144b | 0.44671 | 12 | 0.00049 | 0 | 0.00098 | True |
| *Bacillus pumilus* 145 | 0.49746 | 12 | 0.00049 | 0 | 0.00098 | True |
| *Bacillus pumilus* 144a | 0.56441 | 12 | 0.00049 | 0 | 0.00098 | True |
| *Bacillus pumilus* 150a | 0.59973 | 12 | 0.00244 | 3 | 0.00407 | True |
| *Bacillus pumilus* 156 | 0.89069 | 12 | 0.12939 | 19 | 0.12939 | False |
| *Bacillus subtilis* PY79 | 0.84536 | 12 | 0.01611 | 9 | 0.02302 | True |
| *Bacillus swezeyi* 21 | 0.87747 | 12 | 0.10986 | 18 | 0.12939 | False |
| *Bacillus atrophaeus* 90 | 0.95457 | 12 | 0.12939 | 19 | 0.12939 | False |

**Supplementary Table 6 | Pairwise interaction outcomes between sensitive and resistant strains.** Survival fractions of sensitive targets following co-culture with candidate resistant strains. Data are presented as mean ± s.d. with 95% confidence intervals (CI) from three independent biological replicates (n = 3). Statistical significance was assessed against a neutral monoculture baseline (survival fraction = 1.0) using two-sided one-sample t-tests. P values were adjusted for multiple comparisons using the Benjamini–Hochberg procedure, with significance determined by the resulting adjusted P values (q). Note: Bacillus thuringiensis 111 is abbreviated as Bc111 in the main text to maintain consistency with prior literature.

| Antagonist | Sensitive | N | Mean | SD | CI95_low | CI95_high | p_raw | p_adj | Significance | Mean ± SD (CI95%) |
|---|---|---|---|---|---|---|---|---|---|---|
| *Bacillus thuringiensis* 111 | *Rossellomorea aquimaris* 87b | 3 | 0.873 | 0.063 | 0.801 | 0.944 | 0.073 | 0.137 | | 0.873 ± 0.063 |

| | | | | | | | | | | |
|---|---|---|---|---|---|---|---|---|---|---|
| | | | | | | | | | | (0.801–0.944) |
| *Bacillus thuringiensis* 111 | *Rossellomorea vietnamensis* 451a | 3 | 1.048 | 0.203 | 0.818 | 1.278 | 0.721 | 0.777 | | 1.048 ± 0.203 (0.818–1.278) |
| *Bacillus thuringiensis* 111 | *Sutcliffiella horikoshii* 20a | 3 | 1.009 | 0.054 | 0.948 | 1.069 | 0.801 | 0.831 | | 1.009 ± 0.054 (0.948–1.069) |
| *Bacillus thuringiensis* 111 | *Sutcliffiella horikoshii* 98b | 3 | 0.908 | 0.184 | 0.700 | 1.116 | 0.478 | 0.546 | | 0.908 ± 0.184 (0.700–1.116) |
| *Bacillus cereus* 102 | *Rossellomorea aquimaris* 87b | 3 | 0.868 | 0.122 | 0.730 | 1.006 | 0.202 | 0.290 | | 0.868 ± 0.122 (0.730–1.006) |
| *Bacillus cereus* 102 | *Rossellomorea vietnamensis* 451a | 3 | 0.919 | 0.128 | 0.774 | 1.063 | 0.385 | 0.469 | | 0.919 ± 0.128 (0.774–1.063) |
| *Bacillus cereus* 102 | *Sutcliffiella horikoshii* 20a | 3 | 0.913 | 0.032 | 0.878 | 0.949 | 0.042 | 0.097 | | 0.913 ± 0.032 (0.878–0.949) |
| *Bacillus cereus* 102 | *Sutcliffiella horikoshii* 98b | 3 | 0.853 | 0.095 | 0.745 | 0.960 | 0.115 | 0.197 | | 0.853 ± 0.095 (0.745–0.960) |
| *Bacillus wiedmanii 1*40a | *Rossellomorea aquimaris* 87b | 3 | 0.891 | 0.071 | 0.811 | 0.971 | 0.116 | 0.197 | | 0.891 ± 0.071 (0.811–0.971) |
| *Bacillus wiedmanii 1*40a | *Rossellomorea vietnamensis* 451a | 3 | 1.034 | 0.184 | 0.826 | 1.242 | 0.780 | 0.824 | | 1.034 ± 0.184 |

| | | | | | | | | | | |
|---|---|---|---|---|---|---|---|---|---|---|
| | | | | | | | | | | (0.826–1.242) |
| *Bacillus wiedmanii* 140a | *Sutcliffiella horikoshii* 20a | 3 | 0.804 | 0.090 | 0.702 | 0.906 | 0.064 | 0.133 | | 0.804 ± 0.090 (0.702–0.906) |
| *Bacillus wiedmanii* 140a | *Sutcliffiella horikoshii* 98b | 3 | 0.848 | 0.126 | 0.705 | 0.990 | 0.171 | 0.266 | | 0.848 ± 0.126 (0.705–0.990) |
| *Staphylococcus arlettae* 99b | *Rossellomorea aquimaris* 87b | 3 | 0.877 | 0.160 | 0.697 | 1.058 | 0.315 | 0.410 | | 0.877 ± 0.160 (0.697–1.058) |
| *Staphylococcus arlettae* 99b | *Rossellomorea vietnamensis* 451a | 3 | 1.036 | 0.030 | 1.001 | 1.070 | 0.180 | 0.273 | | 1.036 ± 0.030 (1.001–1.070) |
| *Staphylococcus arlettae* 99b | *Sutcliffiella horikoshii* 20a | 3 | 0.801 | 0.016 | 0.782 | 0.819 | 0.002 | 0.013 | * | 0.801 ± 0.016 (0.782–0.819) |
| *Staphylococcus arlettae* 99b | *Sutcliffiella horikoshii* 98b | 3 | 0.891 | 0.148 | 0.724 | 1.058 | 0.331 | 0.421 | | 0.891 ± 0.148 (0.724–1.058) |

**Supplementary Table 7 | Quantification of higher-order interactions across 34 microbial triads using an additive null model.** Statistical validation of emergent and suppressive community outcomes. The predicted baseline under the additive assumption equals the survival fraction of the sensitive strain (S) interacting solely with the antagonist (A), denoted as f(A,S). *Observed S_survival* represents the survival of S in the full three-species community. Deviations from this baseline identify higher-order interactions (HOIs), with negative values indicating suppressive HOIs and positive values indicating emergent rescue. Statistical significance was assessed using two-sided two-sample *t*-

tests (p2s). *P* values were adjusted for multiple comparisons across all 34 triads using the Benjamini–Hochberg procedure (q2s). E, normalized emergence ratio.

| **Antagonistic strain** | **Resistant Strain** | **Sensitive strain** | ***f(A,S)* (95% CI)** | **Predicted** | **Observed S_survival (95% CI)** | ***f(A, R, S)* (95% CI)** | ***E*** | ***p2s*** | ***q2s*** | **Interpretation** |
|---|---|---|---|---|---|---|---|---|---|---|
| Bp145 | Bc111 | Sh20a | 0.084 (0.010–0.159) | 0.084 | 0.532 (0.241–0.824) | 0.448 (0.254–0.642) | 0.49 | 0.003 | 0.020 | Emergent |
| Bp145 | Bc102 | Sh20a | 0.084 (0.010–0.159) | 0.084 | 0.544 (0.162–0.925) | 0.460 (0.209–0.710) | 0.50 | 0.007 | 0.022 | Emergent |
| Bp145 | Bw140a | Sh20a | 0.084 (0.010–0.159) | 0.084 | 0.190 (0.168–0.212) | 0.106 (0.055–0.156) | 0.12 | 0.004 | 0.020 | Emergent |
| Bp145 | Sa99b | Sh20a | 0.084 (0.010–0.159) | 0.084 | 0.201 (0.084–0.318) | 0.117 (0.027–0.206) | 0.13 | 0.022 | 0.044 | Emergent |
| Bp95a | Bc111 | Sh20a | 0.053 (-0.104–0.210) | 0.053 | 0.157 (-0.007–0.320) | 0.104 (-0.042–0.251) | 0.11 | 0.119 | 0.169 | None |
| Bp95a | Bc102 | Sh20a | 0.053 (-0.104–0.210) | 0.053 | 0.073 (-0.005–0.151) | 0.020 (-0.093–0.133) | 0.02 | 0.646 | 0.758 | None |
| Bp112a | Bc111 | Sh20a | 0.166 (0.071–0.261) | 0.166 | 0.228 (0.104–0.352) | 0.062 (-0.039–0.163) | 0.07 | 0.163 | 0.216 | None |
| Bp112a | Bc102 | Sh20a | 0.166 (0.071–0.261) | 0.166 | 0.269 (0.217–0.320) | 0.103 (0.033–0.172) | 0.12 | 0.015 | 0.034 | Emergent |
| Bp144b | Bc111 | Sh20a | 0.093 (-0.015–0.200) | 0.093 | 0.457 (0.300–0.613) | 0.364 (0.241–0.486) | 0.40 | 0.001 | 0.020 | Emergent |
| Bp144b | Bc102 | Sh20a | 0.093 (-0.015–0.200) | 0.093 | 0.405 (0.194–0.616) | 0.312 (0.159–0.465) | 0.34 | 0.005 | 0.020 | Emergent |

| | | | | | | | | | | |
|---|---|---|---|---|---|---|---|---|---|---|
| Bp145 | Bc111 | Sh98b | 0.000 (0.000–0.000) | 0.000 | 0.000 (0.000–0.000) | 0.000 (0.000–0.000) | 0.00 | 1.000 | 1.000 | None |
| Bp145 | Bc102 | Sh98b | 0.000 (0.000–0.000) | 0.000 | 0.001 (-0.002–0.003) | 0.001 (-0.001–0.002) | 0.00 | 0.374 | 0.471 | None |
| Bp95a | Bc111 | Sh98b | 0.064 (0.053–0.075) | 0.064 | 0.590 (0.167–1.012) | 0.526 (0.253–0.798) | 0.56 | 0.006 | 0.022 | Emergent |
| Bp95a | Bc102 | Sh98b | 0.064 (0.053–0.075) | 0.064 | 0.219 (-0.080–0.519) | 0.155 (-0.038–0.349) | 0.17 | 0.090 | 0.133 | None |
| Bp112a | Bc111 | Sh98b | 0.014 (-0.018–0.045) | 0.014 | 0.051 (0.025–0.076) | 0.037 (0.011–0.064) | 0.04 | 0.017 | 0.037 | Emergent |
| Bp112a | Bc102 | Sh98b | 0.014 (-0.018–0.045) | 0.014 | 0.063 (-0.024–0.150) | 0.049 (-0.010–0.109) | 0.05 | 0.084 | 0.129 | None |
| Bp144b | Bc111 | Sh98b | 0.400 (0.222–0.579) | 0.400 | 0.125 (0.039–0.211) | -0.275 (-0.403–-0.147) | -0.46 | 0.004 | 0.020 | Suppressive |
| Bp144b | Bc102 | Sh98b | 0.400 (0.222–0.579) | 0.400 | 0.413 (0.175–0.650) | 0.012 (-0.179–0.204) | 0.02 | 0.866 | 0.921 | None |
| Bp145 | Bc111 | Rv451a | 0.214 (0.118–0.310) | 0.214 | 0.288 (0.126–0.451) | 0.074 (-0.047–0.196) | 0.09 | 0.165 | 0.216 | None |
| Bp145 | Bc102 | Rv451a | 0.214 (0.118–0.310) | 0.214 | 0.238 (0.163–0.312) | 0.023 (-0.055–0.102) | 0.03 | 0.453 | 0.550 | None |
| Bp95a | Bc111 | Rv451a | 0.217 (0.138–0.297) | 0.217 | 0.010 (-0.011–0.031) | -0.207 (-0.261–-0.154) | -0.26 | 0.000 | 0.014 | Suppressive |
| Bp95a | Bc102 | Rv451a | 0.217 (0.138–0.297) | 0.217 | 0.118 (0.009–0.227) | -0.100 (-0.187–-0.012) | -0.13 | 0.034 | 0.060 | None |

| | | | | | | | | | | |
|---|---|---|---|---|---|---|---|---|---|---|
| Bp112a | Bc111 | Rv451a | 0.207 (0.142–0.272) | 0.207 | 0.216 (0.119–0.312) | 0.009 (-0.066–0.084) | 0.01 | 0.764 | 0.866 | None |
| Bp112a | Bc102 | Rv451a | 0.207 (0.142–0.272) | 0.207 | 0.122 (0.075–0.169) | -0.085 (-0.137–-0.033) | -0.11 | 0.011 | 0.028 | Suppressive |
| Bp144b | Bc111 | Rv451a | 0.229 (0.145–0.313) | 0.229 | 0.223 (0.140–0.306) | -0.006 (-0.082–0.070) | -0.01 | 0.838 | 0.920 | None |
| Bp144b | Bc102 | Rv451a | 0.229 (0.145–0.313) | 0.229 | 0.234 (0.057–0.412) | 0.005 (-0.122–0.132) | 0.01 | 0.918 | 0.945 | None |
| Bp145 | Bc111 | Ra87b | 0.047 (0.014–0.079) | 0.047 | 0.169 (0.062–0.276) | 0.122 (0.050–0.195) | 0.13 | 0.009 | 0.026 | Emergent |
| Bp145 | Bc102 | Ra87b | 0.047 (0.014–0.079) | 0.047 | 0.091 (0.017–0.165) | 0.044 (-0.008–0.096) | 0.05 | 0.078 | 0.126 | None |
| Bp95a | Bc111 | Ra87b | 0.336 (0.120–0.551) | 0.336 | 0.735 (0.563–0.906) | 0.399 (0.221–0.577) | 0.60 | 0.003 | 0.020 | Emergent |
| Bp95a | Bc102 | Ra87b | 0.336 (0.120–0.551) | 0.336 | 0.647 (0.427–0.867) | 0.311 (0.112–0.510) | 0.47 | 0.012 | 0.030 | Emergent |
| Bp112a | Bc111 | Ra87b | 0.230 (0.148–0.313) | 0.230 | 0.418 (0.141–0.695) | 0.188 (0.001–0.374) | 0.24 | 0.049 | 0.083 | None |
| Bp112a | Bc102 | Ra87b | 0.230 (0.148–0.313) | 0.230 | 0.385 (0.218–0.551) | 0.154 (0.034–0.274) | 0.20 | 0.023 | 0.044 | Emergent |
| Bp144b | Bc111 | Ra87b | 0.437 (0.213–0.661) | 0.437 | 0.134 (0.024–0.244) | -0.303 (-0.464–-0.143) | -0.54 | 0.006 | 0.022 | Suppressive |
| Bp144b | Bc102 | Ra87b | 0.437 (0.213–0.661) | 0.437 | 0.067 (-0.063–0.196) | -0.371 (-0.538–-0.204) | -0.66 | 0.004 | 0.020 | Suppressive |

**Supplementary Table 8 | Quantification of higher-order interactions using a multiplicative null model.** Recalculation of predicted triad survival under the multiplicative assumption, where the baseline equals f(A,S) × f(R,S). Empirical survival fractions from pairwise assays were used for f(R,S). The symbol † indicates instances where f(R,S) was strictly set to 1.0 because the pairwise R × S survival did not significantly differ from the neutral baseline after Benjamini–Hochberg correction (q ≥ 0.05). Statistical significance was assessed using two-sided two-sample *t*-tests (p2s). *P* values were adjusted for multiple comparisons across all 34 triads using the Benjamini–Hochberg procedure (q2s).

| **Antagonistic strain** | **Resistant Strain** | **Sensitive strain** | ***f(A,S)* (95% CI)** | ***f(R,S)* (95% CI)** | **Predicted** | **Observed S_survival (95% CI)** | ***f(A, R, S)* (95% CI)** | ***E*** | ***p2s*** | ***q2s*** | **Interpretation** |
|---|---|---|---|---|---|---|---|---|---|---|---|
| Bp145 | Bc111 | Sh20a | 0.084 (0.010–0.159) | 1.000† | 0.084 | 0.532 (0.241–0.824) | 0.448 (0.254–0.642) | 0.49 | 0.003 | 0.020 | Emergent |
| Bp145 | Bc102 | Sh20a | 0.084 (0.010–0.159) | 1.000† | 0.084 | 0.544 (0.162–0.925) | 0.460 (0.209–0.710) | 0.50 | 0.007 | 0.022 | Emergent |
| Bp145 | Bw140a | Sh20a | 0.084 (0.010–0.159) | 1.000† | 0.084 | 0.190 (0.168–0.212) | 0.106 (0.055–0.156) | 0.12 | 0.004 | 0.020 | Emergent |
| Bp145 | Sa99b | Sh20a | 0.084 (0.010–0.159) | 0.801 (0.760–0.841) | 0.067 | 0.201 (0.084–0.318) | 0.133 (0.049–0.218) | 0.14 | 0.012 | 0.028 | Emergent |
| Bp95a | Bc111 | Sh20a | 0.053 (-0.104–0.210) | 1.000† | 0.053 | 0.157 (-0.007–0.320) | 0.104 (-0.042–0.251) | 0.11 | 0.119 | 0.169 | None |
| Bp95a | Bc102 | Sh20a | 0.053 (-0.104–0.210) | 1.000† | 0.053 | 0.073 (-0.005–0.151) | 0.020 (-0.093–0.133) | 0.02 | 0.646 | 0.758 | None |
| Bp112a | Bc111 | Sh20a | 0.166 (0.071–0.261) | 1.000† | 0.166 | 0.228 (0.104–0.352) | 0.062 (-0.039–0.163) | 0.07 | 0.163 | 0.216 | None |

| | | | | | | | | | | | |
|---|---|---|---|---|---|---|---|---|---|---|---|
| Bp112a | Bc102 | Sh20a | 0.166 (0.071–0.261) | 1.000† | 0.166 | 0.269 (0.217–0.320) | 0.103 (0.033–0.172) | 0.12 | 0.015 | 0.032 | Emergent |
| Bp144b | Bc111 | Sh20a | 0.093 (-0.015–0.200) | 1.000† | 0.093 | 0.457 (0.300–0.613) | 0.364 (0.241–0.486) | 0.40 | 0.001 | 0.020 | Emergent |
| Bp144b | Bc102 | Sh20a | 0.093 (-0.015–0.200) | 1.000† | 0.093 | 0.405 (0.194–0.616) | 0.312 (0.159–0.465) | 0.34 | 0.005 | 0.020 | Emergent |
| Bp145 | Bc111 | Rv451a | 0.214 (0.118–0.310) | 1.000† | 0.214 | 0.288 (0.126–0.451) | 0.074 (-0.047–0.196) | 0.09 | 0.165 | 0.216 | None |
| Bp145 | Bc102 | Rv451a | 0.214 (0.118–0.310) | 1.000† | 0.214 | 0.238 (0.163–0.312) | 0.023 (-0.055–0.102) | 0.03 | 0.453 | 0.550 | None |
| Bp95a | Bc111 | Rv451a | 0.217 (0.138–0.297) | 1.000† | 0.217 | 0.010 (-0.011–0.031) | -0.207 (-0.261–-0.154) | -0.26 | 0.000 | 0.014 | Suppressive |
| Bp95a | Bc102 | Rv451a | 0.217 (0.138–0.297) | 1.000† | 0.217 | 0.118 (0.009–0.227) | -0.100 (-0.187–-0.012) | -0.13 | 0.034 | 0.060 | None |
| Bp112a | Bc111 | Rv451a | 0.207 (0.142–0.272) | 1.000† | 0.207 | 0.216 (0.119–0.312) | 0.009 (-0.066–0.084) | 0.01 | 0.764 | 0.866 | None |
| Bp112a | Bc102 | Rv451a | 0.207 (0.142–0.272) | 1.000† | 0.207 | 0.122 (0.075–0.169) | -0.085 (-0.137–-0.033) | -0.11 | 0.011 | 0.028 | Suppressive |
| Bp144b | Bc111 | Rv451a | 0.229 (0.145–0.313) | 1.000† | 0.229 | 0.223 (0.140–0.306) | -0.006 (-0.082–0.070) | -0.01 | 0.838 | 0.920 | None |
| Bp144b | Bc102 | Rv451a | 0.229 (0.145–0.313) | 1.000† | 0.229 | 0.234 (0.057–0.412) | 0.005 (-0.122–0.132) | 0.01 | 0.918 | 0.945 | None |

| | | | | | | | | | | | |
|---|---|---|---|---|---|---|---|---|---|---|---|
| Bp145 | Bc111 | Ra87b | 0.047 (0.014–0.079) | 1.000† | 0.047 | 0.169 (0.062–0.276) | 0.122 (0.050–0.195) | 0.13 | 0.009 | 0.026 | Emergent |
| Bp145 | Bc102 | Ra87b | 0.047 (0.014–0.079) | 1.000† | 0.047 | 0.091 (0.017–0.165) | 0.044 (-0.008–0.096) | 0.05 | 0.078 | 0.126 | None |
| Bp95a | Bc111 | Ra87b | 0.336 (0.120–0.551) | 1.000† | 0.336 | 0.735 (0.563–0.906) | 0.399 (0.221–0.577) | 0.60 | 0.003 | 0.020 | Emergent |
| Bp95a | Bc102 | Ra87b | 0.336 (0.120–0.551) | 1.000† | 0.336 | 0.647 (0.427–0.867) | 0.311 (0.112–0.510) | 0.47 | 0.012 | 0.028 | Emergent |
| Bp112a | Bc111 | Ra87b | 0.230 (0.148–0.313) | 1.000† | 0.230 | 0.418 (0.141–0.695) | 0.188 (0.001–0.374) | 0.24 | 0.049 | 0.083 | None |
| Bp112a | Bc102 | Ra87b | 0.230 (0.148–0.313) | 1.000† | 0.230 | 0.385 (0.218–0.551) | 0.154 (0.034–0.274) | 0.20 | 0.023 | 0.044 | Emergent |
| Bp144b | Bc111 | Ra87b | 0.437 (0.213–0.661) | 1.000† | 0.437 | 0.134 (0.024–0.244) | -0.303 (-0.464–-0.143) | -0.54 | 0.006 | 0.022 | Suppressive |
| Bp144b | Bc102 | Ra87b | 0.437 (0.213–0.661) | 1.000† | 0.437 | 0.067 (-0.063–0.196) | -0.371 (-0.538–-0.204) | -0.66 | 0.004 | 0.020 | Suppressive |
| Bp145 | Bc111 | Sh98b | 0.000 (0.000–0.000) | 1.000† | 0.000 | 0.000 (0.000–0.000) | 0.000 (0.000–0.000) | 0.00 | 1.000 | 1.000 | None |
| Bp145 | Bc102 | Sh98b | 0.000 (0.000–0.000) | 1.000† | 0.000 | 0.001 (-0.002–0.003) | 0.001 (-0.001–0.002) | 0.00 | 0.374 | 0.471 | None |
| Bp95a | Bc111 | Sh98b | 0.064 (0.053–0.075) | 1.000† | 0.064 | 0.590 (0.167–1.012) | 0.526 (0.253–0.798) | 0.56 | 0.006 | 0.022 | Emergent |

| | | | | | | | | | | | |
|---|---|---|---|---|---|---|---|---|---|---|---|
| Bp95a | Bc102 | Sh98b | 0.064 (0.053–0.075) | 1.000† | 0.064 | 0.219 (-0.080–0.519) | 0.155 (-0.038–0.349) | 0.17 | 0.090 | 0.133 | None |
| Bp112a | Bc111 | Sh98b | 0.014 (-0.018–0.045) | 1.000† | 0.014 | 0.051 (0.025–0.076) | 0.037 (0.011–0.064) | 0.04 | 0.017 | 0.035 | Emergent |
| Bp112a | Bc102 | Sh98b | 0.014 (-0.018–0.045) | 1.000† | 0.014 | 0.063 (-0.024–0.150) | 0.049 (-0.010–0.109) | 0.05 | 0.084 | 0.129 | None |
| Bp144b | Bc111 | Sh98b | 0.400 (0.222–0.579) | 1.000† | 0.400 | 0.125 (0.039–0.211) | -0.275 (-0.403–-0.147) | -0.46 | 0.004 | 0.020 | Suppressive |
| Bp144b | Bc102 | Sh98b | 0.400 (0.222–0.579) | 1.000† | 0.400 | 0.413 (0.175–0.650) | 0.012 (-0.179–0.204) | 0.02 | 0.866 | 0.921 | None |

**Supplementary Table 9 | Classification concordance between additive and multiplicative null models.** No triad changed classification between the additive and multiplicative models. For the single combination with an empirically validated *f*(R,S) ≠ 1.0 (Sa99b × Sh20a; *q* = 0.036), the multiplicative correction increased the estimated higher-order interaction (HOI) term by 0.016, reinforcing rather than undermining the emergent classification.

| **Triad** | **Additive *f(A, R, S)*** | **Mult. *f(A, R, S)*** | **Additive class.** | **Mult. class.** | **Changed?** |
|---|---|---|---|---|---|
| Bp145 × Sh20a × Sa99b | 0.117 | 0.133 | Emergent | Emergent | No |
| All other 33 triads | — | — | (Identical to additive) | (Identical) | No |
